# Structural and biophysical characterization of the tandem substrate-binding domains of the ABC importer GlnPQ

**DOI:** 10.1101/2020.12.19.423572

**Authors:** Evelyn Ploetz, Gea K. Schuurman-Wolters, Niels Zijlstra, Amarins W. Jager, Douglas A. Griffith, Albert Guskov, Giorgos Gouridis, Bert Poolman, Thorben Cordes

## Abstract

The ATP-binding cassette transporter GlnPQ is an essential uptake system that transports glutamine, glutamic acid, and asparagine in Gram-positive bacteria. It features two extracytoplasmic substrate-binding domains (SBDs) that are linked in tandem to the transmembrane domain of the transporter. The two SBDs differ in their ligand specificities, binding affinities and their distance to the transmembrane domain. Here, we elucidate the effects of the tandem arrangement of the domains on the biochemical, biophysical and structural properties of the protein. For this, we determined the crystal structure of the ligand-free tandem SBD1-2 protein from *L. lactis* in the absence of the transporter and compared the tandem to the isolated SBDs. We also used isothermal titration calorimetry to determine the ligand-binding affinity of the SBDs and single-molecule Förster-resonance energy transfer (smFRET) to relate ligand binding to conformational changes in each of the domains of the tandem. We show that substrate binding and conformational changes are not notably affected by the presence of the adjoining domain in the wild-type protein, and changes only occur when the linker between the domains is shortened. In a proof-of-concept experiment, we combine smFRET with protein-induced fluorescence enhancement and show that a decrease in SBD linker length is observed as a linear increase in donor-brightness for SBD2 while we can still monitor the conformational states (open/closed) of SBD1. These results demonstrate the feasibility of PIFE-FRET to monitor protein-protein interactions and conformational states simultaneously.

**HIGHLIGHTS:**

- Resolved crystal structure of tandem SBD1-2 of GlnPQ from *Lactococcus lactis*
- Conformational states and ligand binding affinities of individual domains SBD1 and SBD2 are similar to tandem SBD1-2
- No cooperative effects are seen for different ligands for SBDs in the tandem
- Proof of concept experiments show that PIFE-FRET can monitor SBD conformations and protein-protein interaction simultaneously

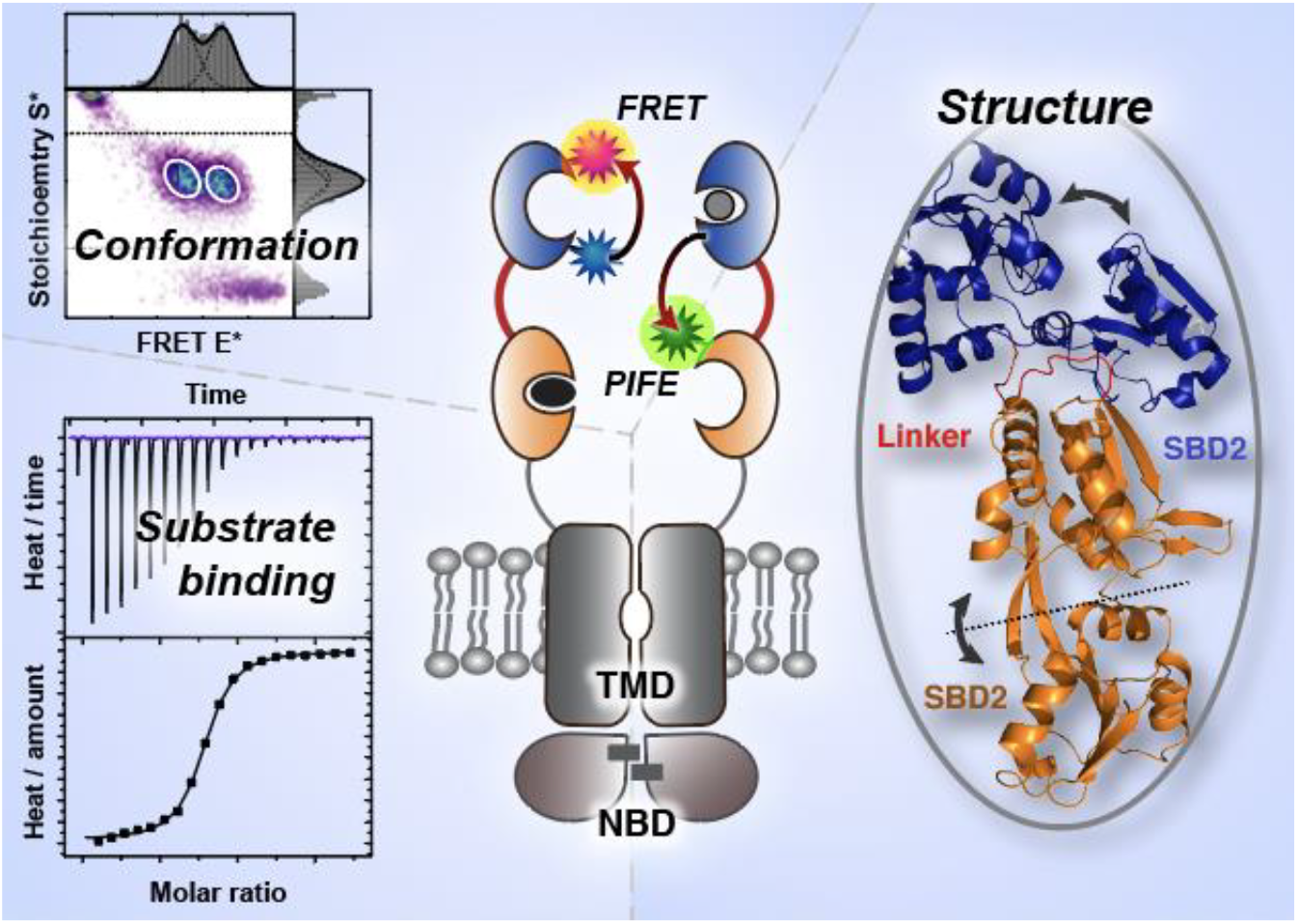

## INTRODUCTION

ATP-binding cassette (ABC) transporters represent a major family of transmembrane proteins, involved in a variety of cellular processes [1–3], including nutrient uptake, antibiotic and drug-resistance, lipid trafficking, and cell volume regulation. They mediate uphill transport of solutes across cellular or organellar membranes using hydrolysis of cytosolic ATP. The core of an ABC transport system is composed of two transmembrane domains (TMDs) and two highly conserved nucleotide-binding domains (NBDs) [4], Figure 1A. In bacterial ABC importers, additional substrate-binding proteins (SBPs) or domains (SBDs) specifically capture and deliver substrates to the TMDs for transport [5, 6]. In some cases, multiple distinct SBPs enable the transport of distinct substrates via the same translocator domain [7–10].

**Figure 1.**
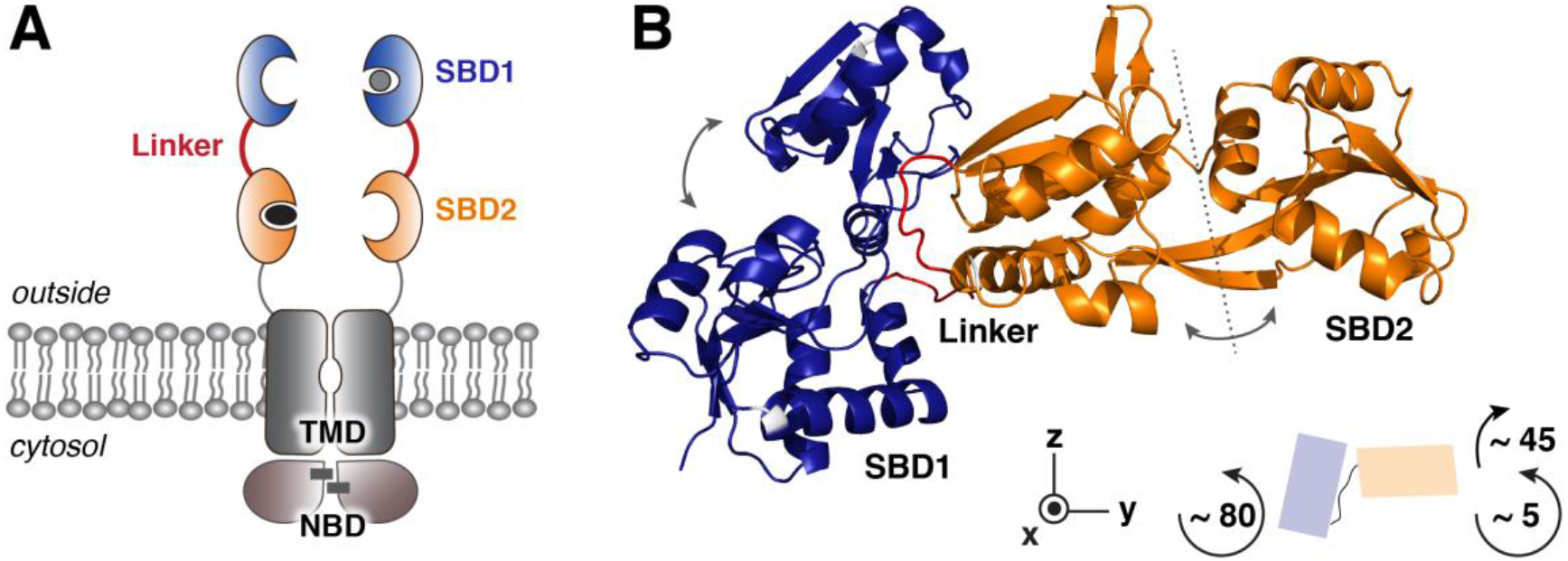
Domain organization of GlnPQ. **(A)** Schematic depiction of GlnPQ. The homodimer is composed of two GlnP and two GlnQ subunits. SBD1 and SBD2 are shown in blue and orange, respectively. Substrates are depicted in grey (asparagine) and black (glutamine). Black bars in the NBDs indicate two molecules of ATP. **(B)** Crystal structure of the tandem SBD1-2 with the same colouring scheme as in A. Both SBDs capture amino acids between their two lobes by closing perpendicularly to their hinge region (grey dashed line; grey arrows). The linker region (red) is close to the hinge region of SBD1. Both SBDs are oriented such that SBD2 appears rotated by ~75° and 45° along the x- and y-axis, respectively. For simplicity only one of the two orientations observed in the crystal structure (chain A) of the SBDs in the tandems is shown.

The ABC import systems are divided into three categories according to the overall structure of their TMDs, which dictates the mechanisms by which they facilitate transport [11–13]. Type I and II ABC importers make use of extra-cytoplasmic SBPs or SBDs that capture ligands directly from the surrounding medium. Substrate uptake is then a multistep process: after binding of the ligand to the SBP, the latter docks onto the TMDs and releases the substrate into a pocket in the translocation pathway. Uphill substrate transport is facilitated by alternating access of the pocket to the opposing sides of the membrane. Type III import systems [14–16] feature a membrane-integrated substrate-binding protein (S factor). The SBPs of Type I/II ABC importers in Gram-negative bacteria are found in the periplasm, where they freely diffuse to capture their substrates. In Gram-positive bacteria, archaea and some Gram-negative bacteria, the SBPs are directly linked to the membrane by a lipid anchor or are tethered to the translocator (hence the name SBD).

In this contribution, we focus on the Type I ABC importer GlnPQ, which features two distinct SBDs in tandem: SBD1 and SBD2 (Figure 1A). This multi-subunit protein is an essential uptake system for glutamine and/or glutamic acid in a variety of non-pathogenic (e.g., *L. lactis*) and pathogenic Gram-positive bacteria (e.g., *S. pyogenes, S. aureus, E. faecalis*), but the systems can also transport a variety of non-essential amino acids [7–10]. GlnPQ is a homodimer composed of two subunits: GlnP and GlnQ. GlnP comprises the TMDs, which are C-terminally linked to SBD2 and SBD1, whereas GlnQ is the cytosolic NBD. SBD2 is attached to the TMD via a 19 amino-acid long flexible linker (Figure 1A; grey) and connected to SBD1 via a 14 amino-acid long linker [17] (Figure 1A, red). This arrangement facilitates the fast delivery of ligands from the SBDs to the translocator. GlnPQ imports glutamine, glutamic acid and asparagine [8]. Whereas the proximal SBD2 exclusively binds glutamine with a K_D_ of ~ 0.9 μM, the distal SBD1 binds both: asparagine with high affinity (K_D_ = 200 nM) and glutamine with a low affinity (K_D_ = 90 μM) [8]. The SBDs have thus evolved distinct substrate specificity.

Both crystallography and single-molecule Förster-resonance energy transfer (smFRET) experiments on the single SBD1 and SBD2 showed that substrate binding was linked to a conformational change of the corresponding SBD from an open *apo* conformation to a closed liganded conformation [18–21]. The results also implied an induced-fit-type ligand-binding mechanism, where conformational dynamics are induced by ligand-SBD interactions similar as later also demonstrated for other SBPs [22–26]. Additionally, it was shown that the opening of the SBDs and ligand release was the rate-limiting step in the transport cycle and that the closed conformation triggers ATP-hydrolysis and transport [18]. More recently, it was shown that some SBPs and SBDs can recognize multiple distinct ligands and that the ligand-SBP or SBD complexes formed do not necessarily share a single translocation competent conformation [19]. Instead, transport specificity was determined by the formation of conformers capable of allosteric coupling with the translocator, while retaining conformational dynamics permissive for ligand release. These recent findings provide a new understanding of the mechanistic diversity that enables ABC importers to achieve substrate selectivity [19, 27].

A particularly interesting feature of the GlnPQ importer is the presence of the two SBDs, fused in tandem to the TMD, generating four substrate-binding sites close to the translocation pathway and SBD competition for docking onto the translocator. Even though the mechanism of ligand binding for the individual SBDs is understood, it is not clear how interactions between the SBDs might affect transport. Although we could recently show that changes in the inter-domain distances can affect transport and ATPase activity [17, 28], what this reveals about the native transport mechanism is as yet not fully clear. Also, possible domain interactions or functional cooperativity between the SBDs in the tandem still must be assessed. The key questions are whether the properties of the single SBDs are the same as when they are present in the tandem and whether there is evidence for functional cooperativity, e.g., that binding of substrate to one SBD alters ligand binding or conformational dynamics of the other.

In this work, we therefore focussed on studying the structural and biochemical consequences of connecting two SBDs by a flexible linker. We present crystallographic, biochemical, and biophysical data. We first determined the crystal structure of the SBD-tandem in its ligand-free form and used smFRET-based spectroscopy to determine the underlying conformations and substrate-binding affinities of the individual SBDs within the tandem to disentangle the contributions of the individual domains. We find that tandem ligand-binding domains have identical structures as compared to isolated SBDs and both domains operate largely independently of each other in the tandem. The ligand-binding was only affected marginally by the adjoined domains for extremely short artificial linkers. This finding raises the question about how optimized the length of the linker connecting the two SBDs in GlnPQ is and whether cooperativity can be induced by changing this length. To elucidate the interaction of the two domains and the flexibility provided by the connecting linker in the tandem, we employed inter-domain FRET and explored the suitability of our recently introduced PIFE-FRET assay [29, 30], which combines smFRET with protein-induced fluorescence enhancement (PIFE) for the study of protein-protein interactions.

## RESULTS

### Crystallization and Structure determination

We solved the crystal structure of the unliganded tandem SBD1-2 domain at 2.8 Å resolution (PDB ID 6H30; see Figure 1B, Figure S1 and Table S1). Crystals of the unliganded tandem SBD1-2 in buffer supplemented with MES were grown with the hanging drop vapour diffusion method. The crystals belonged to the C222_1_ space group and contained two polypeptide chains per asymmetric unit with 58% solvent content (Figure S1A). Each of the two chains comprises two SBDs linked via a 14-amino acid loop. The individual SBDs consist of two α/β subdomains. In SBD1, the large α-domain comprises residues 29-113 and 207-251, while the small β-domain is made up of residues 114-206 (see Figure 1B, blue domain). The large α-domain in SBD2 is formed by residues 255-345, and residues 346-440 are of the small β-domain (see Figure 1B, orange domain). Both domains in SBD1 and SBD2 are connected by two anti-parallel β-strands, a common feature in substrate-binding proteins. The two SBDs are structurally classified in the sub-cluster F-IV [5]. The binding site for the substrates is localized between the two domains.

### Structural comparison of single and tandem SBDs

The crystallized tandem SBD1-2 structure reveals MES molecules in the binding pockets of the open state (Figure S1). An asymmetric unit contains two SBD1-2 monomers that are oriented head to tail (Figure S1A). We found that the SBDs in the tandem have identical structures to those of the individual SBDs as revealed by the superposition of SBD1-2 structure with those for unliganded SBD1 (PDB ID 4LA9, rmsd of 0.5 Å) and SBD2 (PDB ID 4KR5, rmsd of 1.1 Å) (Figure S1B; Table S1). The linker sequence (depicted in red in Figure 1 and S1) connects the last α-helix of SBD1 to the first β-sheet of SBD2 and comprises the residues Gly-248 to Val-261. In comparison to other homologs, this sequence is very short [8, 17]: close homologs of SBD1-2 found in *S. pneumoniae* and *E. faecalis* show an extra insertion in this region of 11 amino acids making the linker almost twice as long. We speculate that the connecting linker between the domains should still provide some flexibility as suggested from the way molecules are packed within the crystal (Figure S1C). Both domains of the tandem SBD1-2 are oriented differently within the crystal unit cell: the superposition along SBD1 of both domains in tandem SBD1-2 reveals a rotation of ~ 45 degrees for SBD2 with close contact to the hinge region of SBD1. For simplicity, only one of the two orientations observed in the crystal structure (chain A) is shown in Figure 1B.

### SBD substrate affinity and specificity

We next analysed the binding properties of the tandem SBD1-2 in comparison to the published ones from single SBD1 and SBD2, using isothermal titration calorimetry (ITC). Figure 2 shows the binding curves for the two high-affinity ligands of SBD1-2 (see Table S2 for details). SBD1 within the tandem SBD1-2 binds asparagine with a dissociation constant K_D_ of 400±100 nM (Figure 2A), which is similar to isolated SBD1 (K_D_ = 200±100 nM; [8]). The titration of SBD1-2 with glutamine reveals two binding sites, one with high and one with low affinity since glutamine can be bound by both SBDs (Figure 2B). The K_D_ values for binding of glutamine were 0.6±0.2 μM (for SBD2) and 180±100 μM (for SBD1), which are similar to the values observed for the isolated SBDs (Table S2 and ref. [8]). In the presence of saturating concentration of asparagine, the K_D_ for binding of glutamine to SBD2 in the tandem and the single domain are the same (Figure 2C). We can thus conclude that the proximity of the domains in the tandem, which is enforced by the linker does not alter ligand affinities.

**Figure 2.**
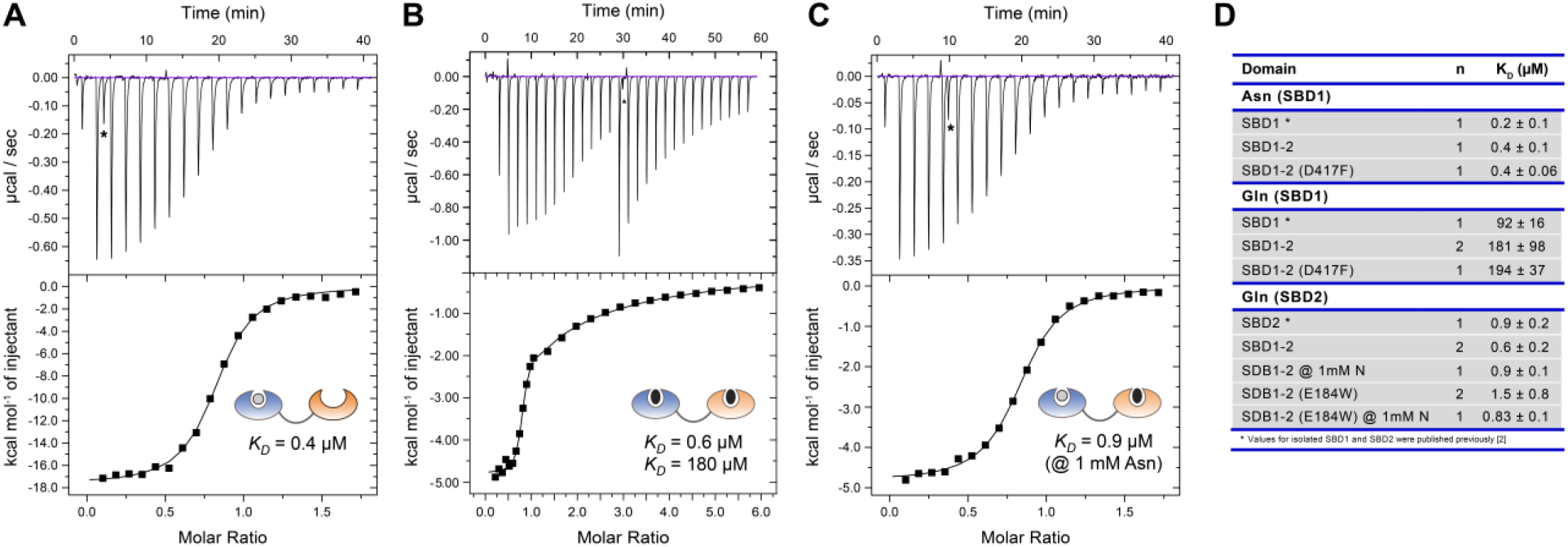
Ligand binding affinities of tandem SDB1-2 as determined by ITC. **(A)** Binding of asparagine to SBD1-2 occurs to SBD1, shown in blue. **(B)** In the absence of asparagine, glutamine binds to both domains in the tandem with KD values of 180 μM and 0.6 μM for SBD1 and SBD2, respectively. **(C)** SBD2 binds glutamine with a KD of 0.9 μM, which was determined by blocking SBD1 with a high concentration (1 mM) of asparagine. Spikes due to leakage of the syringe (marked by *) have been excluded from the analysis. **(D)** Overview of KD values that are summarized in Supplementary Table S2. The KD values were obtained from five biological replicate. Fit values for KD are based on the data shown in the figure.

This conclusion is further supported by experiments on mutants with one inactive and one functional ligand-binding domain. It was shown previously that isolated variants SBD1(E184W) and SBD2(D417F) do not bind ligand and remain in the open conformation even in the presence of substrates [18, 31]. ITC experiments on similar tandem variants SBD1(E184W)-2 (inactive SBD1) and SBD1-2(D417F) (inactive SBD2) show that the functional SBD of the tandem is not affected by the inactivation of the other SBD (Figure S2, Table S2). When comparing SBD1(E184W)-2 to the wildtype SBD1-2 (Figure S2A), we can conclude that glutamine-induced conformational fluctuations in SBD1 do not affect the binding of glutamine to SBD2. Similarly, experiments with SBD1-2(D417F) (Figure S2B-C) imply that the binding of a ligand to the SBD2 does not affect binding to SBD1. In accordance, the binding isotherms of SBD1-2 for glutamine are a superimposition of those of SDB1 plus SBD2.

### Binding affinities and states of single and tandem-SBDs probed by smFRET

In addition to our ITC experiments, we also used smFRET assays as an independent approach to examine whether there is functional cooperativity between the domains in the tandem. We employed smFRET [32–35] in a fashion similar to previous work [18, 19] to monitor conformational state changes and to simultaneously extract the substrate binding affinity of the individual domains within the tandem. In the smFRET assay (Figure 3A), we observe conformational changes directly as differences in the FRET efficiency, where the open conformation is characterized by a low FRET state and the closed substrate-bound conformation has a higher FRET state (Figure 3B). We employed different cysteine variants as described previously [18, 19] and created the corresponding variants for the tandem SBD1-2 (Figure 3A). Cysteine residues were located at G87C and T159C in SBD1 of the tandem SBD1-2 (Figure 3A, i-ii) and T369C and S451C in SBD2 of SBD1-2 (Figure 3A, iii-iv). We denote the two cysteine-backgrounds as subscripts on the single SBDs, such as SBD1^A^-2 and SBD1-2^C^(a summary of the short notations of all proteins including mutations is given in Table S3). The occurrence of potentially problematic fluorophore-protein interactions were ruled out by steady-state anisotropy experiments (Table S4).

**Figure 3.**
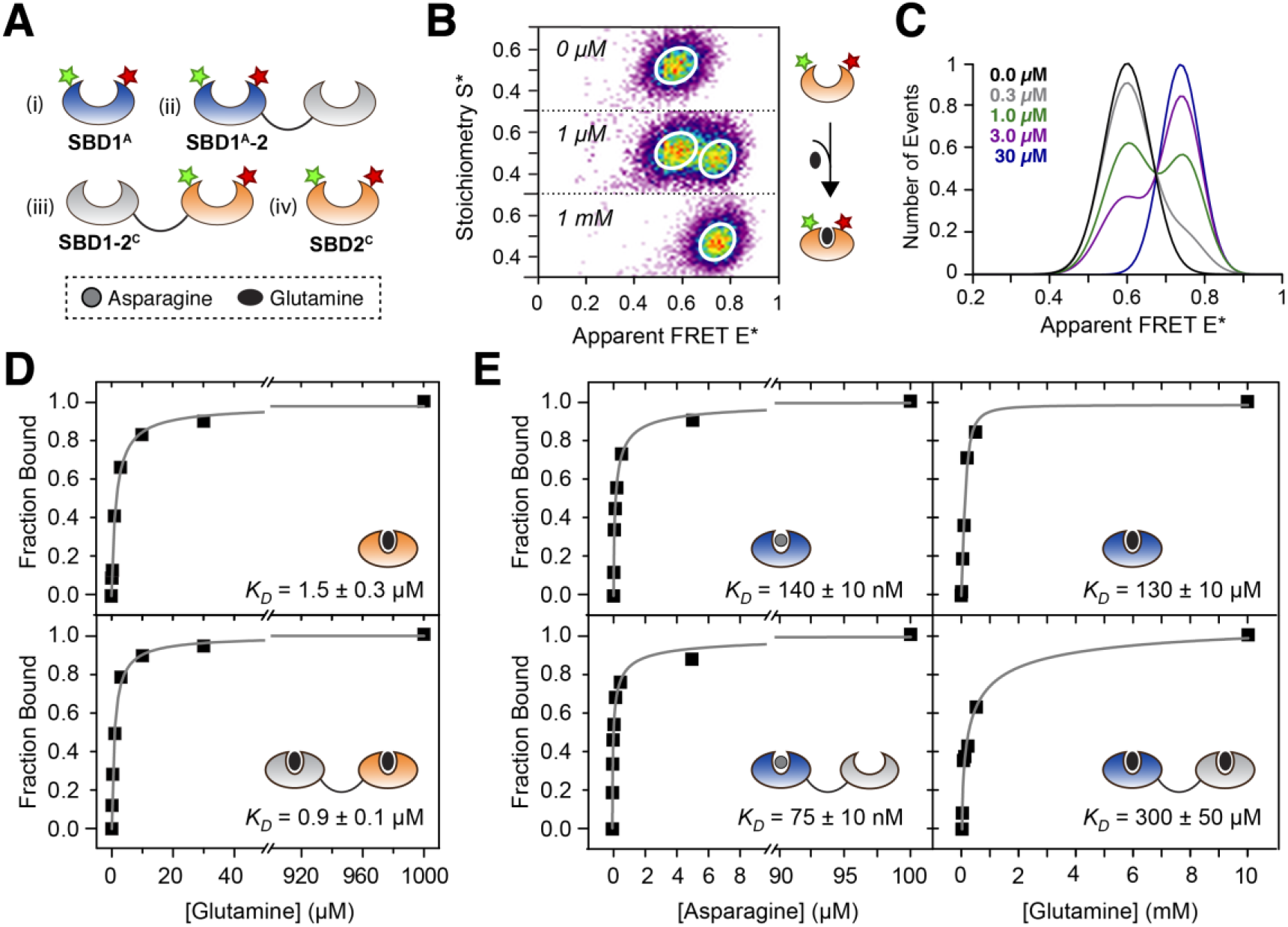
ALEX spectroscopy on single and tandem SBDs in GlnPQ. **(A)** Design of smFRET assay to monitor intramolecular SBD conformational states. **(B)** Confocal based ALEX spectroscopy of SBD2^**C**^ labelled with Alexa Fluor 555 and Alexa Fluor 647 maleimide in the presence of 0 μM, 1 μM and 1mM of glutamine. SBD2^**C**^ shows an apparent FRET value of 0.58 in the unliganded state. The apparent FRET E* shifts to 0.74 under saturating concentrations of glutamine. **(C)** Apparent FRET E* histograms as a function of varying ligand concentration. The addition of glutamine to SBD2^**C**^ shifted the population of molecules from a low (unliganded) to high FRET state (closed liganded). At concentrations close to KD both populations were similar. **(D-E)** Binding affinity determination of single and tandem SBDs by ALEX spectroscopy. **(D)** SBD2^**C**^ as single domain and tandem SBD1-2^**C**^ gave KD values for glutamine of 1.5 and 0.9 μM, respectively. The ratio of populations is defined as the ratio of areas in case of the closed liganded population versus (unliganded and closed-liganded population). **(E)** SBD1^**A**^ as single domain and tandem SBD1^**A**^-2 gave an apparent K_D_ for asparagine of 140 and 75 nM, respectively; the corresponding values for glutamine are 130 and 300 μM. Errors indicated were obtained directly from the fit in the respective data set.

We examined ligand binding by stepwise addition of substrate to a very dilute protein solution (≈ 50 pM) and monitored the conformational transition between the open unliganded and the closed-liganded state, which is manifested as a change in FRET efficiency (Figure 3B). We employed μs-ALEX spectroscopy [36–38] with alternating laser excitation at 532 and 640 nm, where fluorescently labelled biomolecules diffuse through the excitation volume of a confocal microscope. After stochastic labelling of SBD2^C^ and the tandem SDB1-2^C^ with Alexa Fluor 555- and Alexa Fluor 647-maleimide, a single population was observed that was distributed around an apparent FRET efficiency E* of 0.58 (Figure 3B; Figure 4A–B; Figure S3 and Table S5–S6). Under saturating concentrations of glutamine, the apparent FRET E* value shifted to 0.74 for both proteins (Figure S3), supporting the idea that both proteins undergo identical conformational changes. Sorting the molecules at the given substrate concentrations according to their FRET value (Figure 3C) revealed that the amplitude of the *apo*-protein gradually decreased with an increasing concentration of ligand, while the closed, liganded state at 0.74 increased in parallel. We obtained the binding curve from the ratio of the number of molecules in the closed-liganded state over the total number of recorded molecules (Figure 3D). It yielded an apparent K_D_ of ~ 0.9 μM for SBD1-2^C^ and ~ 1.5 μM for SBD2^C^, findings that are consistent with ITC experiments (Figure 2 and Table S2). From this we can conclude that glutamine binding to SBD2 was not notably affected by the presence of SBD1 and ligand binding correlates with conformational changes in isolated SBD2 and the tandem.

**Figure 4.**
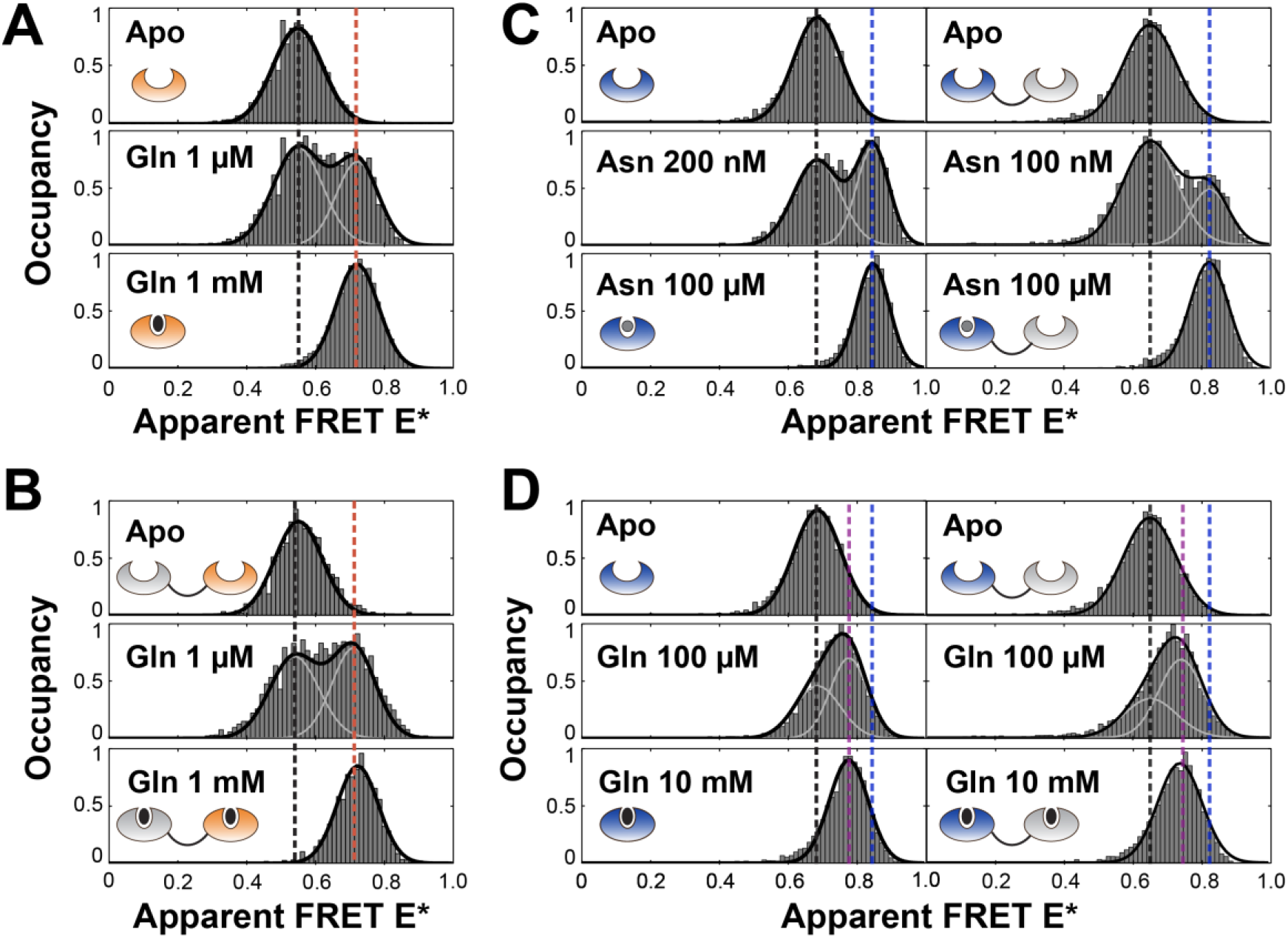
Conformational states of isolated and tandem SBDs probed by ALEX Spectroscopy. **(A-B)** Single and tandem-linked SBD2^**C**^ in the presence of glutamine. Both proteins are characterized by two FRET states: the *apo-*state at 0.58 (black line) and closed-liganded state at 0.74 (orange line). **(C-D)** Single and tandem-linked SBD1^**A**^ mutant in presence of different ligands. Both proteins show a low FRET value of ~ 0.65 in the *apo-*state (black line). **(C)** Upon addition of asparagine: both proteins start closing, which is observed as an additional high FRET state at ~ 0.82 (blue line). At the K_D_, two distinct populations are observed at an equal ratio. For saturating concentrations of asparagine, both mutants are fully closed. **(D)** In presence of glutamine, a gradual shift in FRET is observed hinting towards fast interconversion of states. For saturating concentration, both mutants show an intermediate FRET state of ~ 0.74 (purple line), which is lower than in the case of asparagine (blue line).

In contrast to SBD2, SBD1 binds both asparagine and glutamine (Figure 3E; Figure S4–S5). For smFRET experiments, SBD1 was labelled at G87C and T159C with Alexa Fluor 555 and Alexa Fluor 647 maleimide. The variant showed a single population with an apparent FRET of 0.64 in the *apo*-state (Figure 4C). In the presence of asparagine and glutamine, SBD1^A^-2 undergoes a conformational transition to the same closed liganded state as for isolated SBD1^A^ (Figure 4C/D). The apparent K_D_ of 75 nM for asparagine binding to SBD1^A^-2 (Figure 3E, left) is similar to that obtained for isolated SBD1 (K_D_ = 140 nM). The respective K_D_ values for glutamine binding are also similar with values of 300 μM for tandem SBD1^A^-2 and 130 μM for SBD1^A^.

We conclude from a combined inspection of the ITC and smFRET experiments that (i) the ligand dissociation constants of single and tandem SBDs are not notably different. (ii) Isolated SBDs and their tandem counterparts show identical conformational states in the presence and absence of their ligands, i.e., both high and low-affinity ligands trigger formation of similar ligand-bound closed states.

### Domain orientation of SBD1 and SBD2 in the tandem in solution

To examine whether the flexible linker allows domain re-orientation within the tandem, and to determine the functional relevance of the orientation of the domains in the crystal structure, we employed an inter-domain single-molecule FRET assay. For this we designed a double cysteine variant with one fluorophore anchor point in SBD1 (A136C) and one in SBD2 (T369C), which we denoted as SBD1^T^-2^T^. As a structural reference, we selected chain A from the crystal structure which suggests a ~ 65 Å inter-probe distance for the variant considering the C_β_ distances of the respective amino-acids. We then labelled the variant with Alexa Fluor 555 and Alexa Fluor 647 (Figure 5A). The resulting smFRET histogram in the absence or presence of ligand shows a single population at low apparent FRET efficiency around 0.33 (Figure 5A).

**Figure 5.**
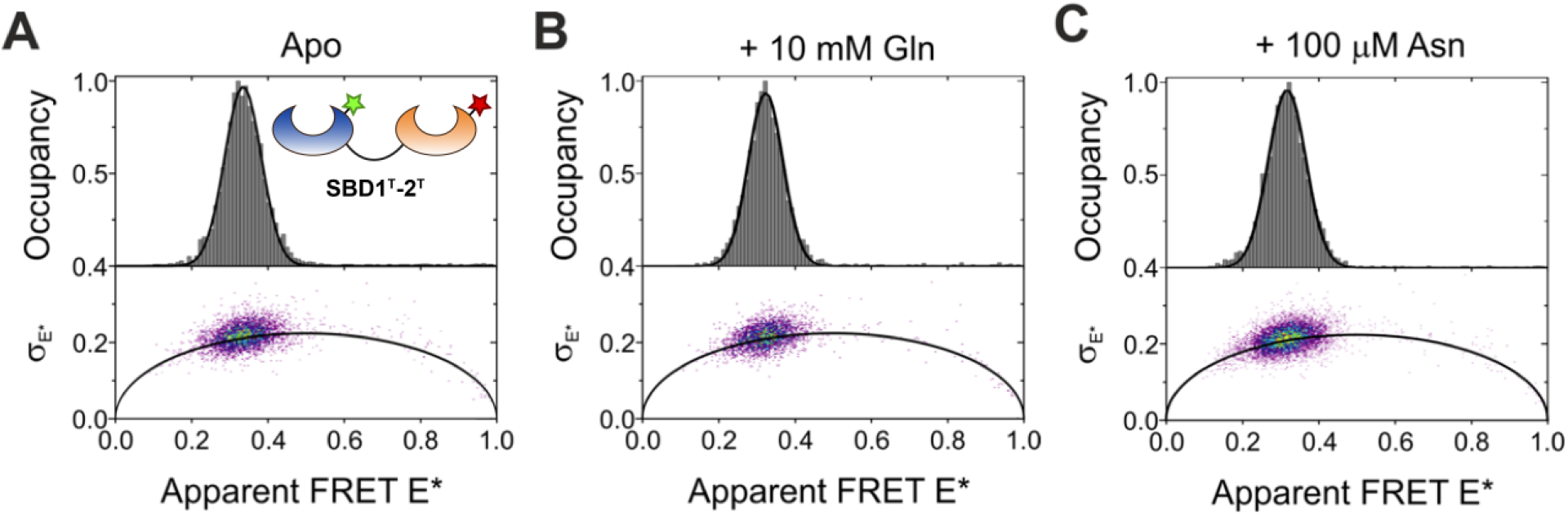
μs-ALEX Spectroscopy and burst variance analysis to probe the inter-domain distance and dynamics in the tandem SBD1^T^-2^T^. **Top:** FRET histograms of the tandem SBD1^T^-2^T^ labelled with Alexa Fluor 555- and Alexa Fluor 647-maleimide, in the **(A)** *apo-*state and under saturating concentrations of **(B)** glutamine and **(C)**. The protein shows a low FRET value of 0.33 in the *apo-*state, which does not change upon addition of ligands; also, the width of the distribution does not change. Double-labelled protein species were identified in the ES-histograms using a stoichiometry range of S = 0.25-0.6.

In addition to the centre position of the FRET distribution, its width also reports on the underlying dynamics in the system. In contrast to the molecules in static samples where any broadening of the peak would exclusively originate from shot-noise, additional fast conformational transitions of the molecules in dynamic samples, on the order of the diffusion time, will result in an additional broadening [39]. A burst variance analysis on the μs-ALEX data of SBD1^T^-2^T^ tandem shows that the width of the population with 0.046 hardly deviates from the theoretical shot-noise limit of 0.037, which was determined using the mean number of photons per burst, which was similar for each condition (Figure 5). These small differences in width implies that the two large domains do not re-arrange their position, or the process is much faster compared to the transit time through the confocal volume. The latter seems more likely since protein rotation should occur on timescales of 10-100 ns (considering the masses of SBD1/2), which is sufficient to allow both SBDs to adapt all relative possible orientations using the linker region as a flexible element. Moreover, the addition of saturating ligand concentrations does not change the position of the peak and thus the distance between the domains, nor the width of the peaks (Figure 5B-C). It is clear from this data set that smFRET will only allow further investigations when combined with PIE-MFD measurements that allow the protein system to be probed on the micro-to nanosecond time scale [40].

### Design and biochemical characterization of linker mutants

To alter the short-distance interactions between both SBDs within the tandem SBD1-2 and to elucidate the effect of the linker length on the properties (ligand affinity and conformational states) of the tandem, we designed a range of SBD1-2 tandems with different linker lengths connecting both SBDs (Figure 6/S6).

**Figure 6.**
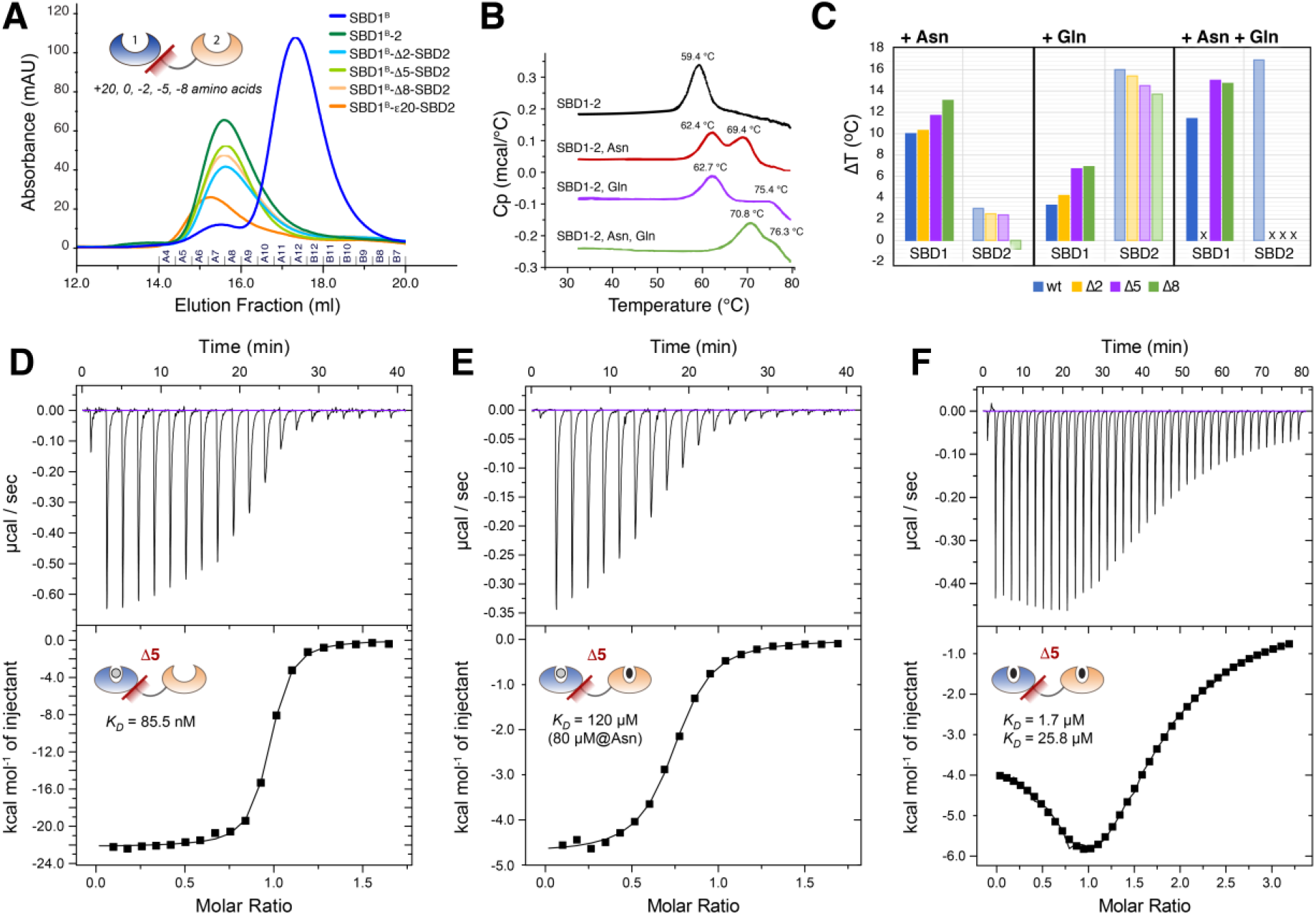
Effects of variations in linker length on the hydrodynamic radius, stability and ligand binding activity of tandem SBD1-2. **(A)** Size-exclusion of unlabelled and cysteine-containing variants on a Superdex 200 10/300 GL column. Tandem SBDs eluted around 15.5 ml and were clearly discernible from the single SBDs, which eluted around 17.5 ml. **(B-C)** Absolute and relative thermostability of SBD1-2 wild-type protein as measured by DSC. **(D-F)** ITC titrations for SBD1-Δ5-SBD2 with **(D)** asparagine, **(E)** glutamine with saturating amounts of asparagine and **(F)** glutamine.

The tandem mutants have deletions of 2, 5 and 8 amino acids within the linker at the C-terminus of Ala-251, and the deletions were made in the cysteine-backgrounds SBD1(A136C/S221C)-SBD2 and SBD1-SBD2(T369C/S451C); see Figure S6A. We denote them as SBD1^(B)^-Δ#-SBD2 and SBD1-Δ#-SBD2^(C)^, wherein # indicates the number of amino acids that are deleted from the linker sequence (Figure S6B). The superscripts refer to the cysteine-background of the mutants. We additionally produced two mutants with an extended linker of 20-amino acid, inserted between Ala-251 and Thr-252. They are named SBD1^B^-ε20-SBD2 and SBD1-ε20-SBD2^C^. A summary of the short notations for all mutants can be found in Table S3. The linker modifications did not alter the apparent hydrodynamic radius of the SBDs compared to the wild-type protein as determined by size-exclusion chromatography (**Figure 6A**), except for SBD1^B^-ε20-SBD2.

Furthermore, we verified the stability of the proteins using differential scanning calorimetry (DSC) by analysing thermostability and potential differences in folding of wild-type SBD1-2 and linker mutants (Figure 6B/C and Table S7). In the unliganded state, the melting temperature of both single SBDs was 59 °C [41], which is the same as that of all tandem SBD1-2 mutants. The addition of either asparagine or glutamine increased the protein stability of the tandem resulting in a higher melting temperature of ~ 62°C for SBD1 and ≥ 69°C for SBD2. SBD1 and SBD2 are unfolding separately as seen by a double peak Figure 6B. We assign the peak shifts to a specific SBD by comparing the melting temperatures of the tandem SBD1-2 to that of the single SBDs [41]. Figure 6C shows the relative thermostability of both domains for all tandem linker mutants in the presence of ligand. We find that linker deletions of 5 and 8 amino acids increase the stability of the proteins by 2-3°C. Maximal stabilization of SBD2 in SBD1-2 tandems is observed in the presence of glutamine, which was increased further in the presence of asparagine (Figure 6C). These data suggest that the SBDs are stabilized by direct protein-protein interactions that do not occur when the linker is too long.

To characterize their biochemical properties, we analysed the linker deletion mutants by ITC and determined the thermodynamic parameters of ligand binding (ΔH, TΔS, ΔG and K_D_; see Table S2). Wild-type SBD1-2 has a single binding site for asparagine and two binding sites for glutamine. The measurements with SBD1-ε20-SBD2 and SBD1-Δ2-SBD2 corroborate our observations that there is no apparent cooperativity in the binding of amino acids by the presence of the adjoining SBD with long linkers. On the other hand, the SBD1-Δ5-SBD2 mutant shows an increased binding affinity for asparagine in SBD1 (**Figure 6D**), but there is only a minor effect on the binding of glutamine to SBD2. The K_D_ for asparagine binding in SBD1 decreases from 0.4 to 0.06 μM for SBD1-Δ5-SBD2. Also, we observe more clearly than in the wild-type protein two glutamine binding sites in SBD1-Δ5-SBD2, indicating that the K_D_ for glutamine binding of SBD1 is also decreased by the deletion of 5 amino (**Figure 6E-F**) from ~ 100 μM to 19 μM (Table S2).

### Simultaneous observation of inter- and intra-domain distances via PIFE-FRET

Next, PIFE-FRET [29, 30], i.e., a synergistic combination of smFRET (Figure A) with protein-induced fluorescence enhancement (PIFE, Figure B), was used to simultaneously study inter- and intra-domain interactions in the tandem SBD1-2 (Figure C). PIFE-FRET was recently introduced by us to monitor interactions between nucleic acids and proteins concomitant to conformational changes [29, 30]. A similar PIFE-FRET assay in μsALEX experiments has, to the best of our knowledge, not been introduced to monitor protein-protein interactions and conformational motion. In the designed experiments, we aimed to probe intradomain distance d_1_ via FRET, while probing the inter-domain dynamics via distance d_2_ using PIFE (Figure C).

PIFE is based on the change of excluded volume or micro-viscosity of an environmentally sensitive dye by the proximity of an adjacent protein and can be observed as a change in fluorescence brightness, lifetime or anisotropy of the dye [29, 30, 42]. To visualize the short distance d_2_ in ALEX experiments, we used the ratiometric brightness ratio S (or stoichiometry), which is defined as the brightness ratio between donor and acceptor fluorophore. A brightness increase of the donor fluorophore due to the proximity of an adjacent protein moiety is thus observed as an increase in the stoichiometry S (Figure C), whereas a brightness increase of the acceptor would lead to a decrease in stoichiometry. We employed the cyanine dye Cy3 as PIFE-sensor in combination with Atto647N as environmentally insensitive FRET acceptor fluorophore. Following our theoretical framework [29, 30], contributions of FRET and PIFE to the emission of the donor and acceptor can be disentangled and both the intra-domain as well as inter-domain distance can be monitored simultaneously. In the case of the tandem SBD1-2 in GlnPQ, PIFE can easily be combined with smFRET, since the assay only requires a specific set of fluorescent labels but is applicable to the same mutants as used for smFRET. The assay has thus the potential to map the interactions between both SBDs (Figure C, Stoichiometry axis) and the conformational states of an individual SBD (Figure C, FRET axis) simultaneously by a mere change of fluorophores.

### PIFE-FRET monitors protein-protein interaction between the two SBDs

To study the interactions between the two SBDs in further detail by PIFE-FRET, we labelled one of the SBDs within the tandem with Cy3/Cy3B- and ATTO647N-maleimide. In this assay, we anticipated a brightness change of Cy3 due to the presence of the adjoining SBD (Figure C). The goal was a comparison of the mean FRET and stoichiometry value of the single SBDs (SBD1^B^ and SBD2^C^), the tandem SBDs (SBD1^B^-2 and SBD1-2^C^), and the linker mutants (SBD1^B^-Δ#-SBD2 and SBD1-Δ#-SBD2^C^), where S-changes would be indicative of PIFE-effects caused by domain-domain interactions. Labelling with Cy3B would serve as a negative control with a donor-fluorophore that does not show PIFE [29, 30].

Based on the crystal-structure of chain A and accessible volume (AV) calculations [43], we built a simple (and maybe oversimplified) model for AV changes caused by the absence and presence of the second SBD (Figure 8A/B). Based on the models we hypothesized that for SBD1^B^-2 tandem and its linker mutants, only conformational changes d_1_ should be observable since no reduction of fluorophore accessible volume is expected in the tandem (Figure 8A). Based on the model in Figure 8B we expect PIFE to occur when Cy3 labels position Ser-451 in SBD2 due to the steric hindrance caused by the neighbouring domain and consecutive reduction of the accessible volume of the dye in this case (Figure 8B). It is important to note that we perform stochastic labelling and that consequently only one of the two resulting sub-populations of labelled SBD2-proteins in the tandem (with the donor located at Ser-451) is expected to show a PIFE-signal. This fact requires careful checking of the observed effects in relation to variations of the donor-acceptor labelling ratio or if available site-specific labelling.

To test our predictions, we investigated single SBD1^B^, SBD1^B^-2 and SBD1^B^-Δ#-SBD2 variants via μs-ALEX and analysis of two-dimensional E/S histograms. The comparison of SBD1^B^ and tandem SBD1^B^-2 showed identical E* distributions with means of 0.56 and 0.63 in the apo and closed liganded states on the X-axis, respectively (Figure C and Figure S7). As observed for SBD1^A^ and SBD1^A^-2, the binding affinity and conformational states in SBD1^B^-2 are not affected by the adjoining SBD2 (Figure S7 and Table S5/S6). Moreover, this holds when shortening the linker between both domains, as shown, e.g., for SBD1^B^-Δ5-SBD2 (Figure S7). Interestingly, there was no significant difference in stoichiometry S* for comparison of single SBD1^B^, SBD1^B^-SBD2 and SBD1^B^-Δ5-SBD2 or others linker variants (Figure 8C/S8), which can be inspected via the mean stoichiometry of each state. This statement holds for the apo state (Figure 8C, green line, S = 0.284) and liganded holo state (Figure 8C, orange line, S = 0.290) for direct comparison of SBD1^B^ and SBD1^B^-Δ5-SBD2, an observation in line with the predictions based on the AV-simulations of chain A (Figure 8A).

Next, we characterized SBD2^C^, SBD1-2^C^ and SBD1-Δ#-SBD2^C^ mutants. Again, the single SDB2^C^ serves as a reference with one population centred at E* = 0.61 (Figure D). Under saturating concentration of glutamine, the apparent FRET species shifts to 0.73. SBD2^C^ and SBD1-2^C^ show the same binding affinity and identical FRET states in the presence and absence of glutamine (Figure S9). However, the presence of the second SBD1 leads to an increase in stoichiometry from 0.291 (apo, single SBD) to 0.306 (apo, tandem SBD) and 0.339 (apo, SBD1-Δ5-SBD2^C^), also for the holo state (Figure D, green line: apo, orange line: holo).

Motivated by the small, yet significant changes, we investigated the systematics of stoichiometry changes in SBD1-Δ#-SBD2^C^ variants as a function of linker length (Figure E/F). Starting from single SBD2, we observe a (linear) increase in stoichiometry for the tandem SBD1-SBD2^C^ reaching a maximum for SBD1-Δ5-SBD2^C^ variant and a minimum value for SBD1-ε20-SBD2^C^. These results are in line with a distance-dependent PIFE effect [29] and reveal the universal nature of PIFE as a molecular ruler, which can be applied even for protein-protein interactions as demonstrated in Figure. To validate the interpretation of S-changes and the observed trends, we also performed labelling with Cy3B as a donor fluorophore, which has identical spectral properties as Cy3, but does not show PIFE effects [29] and can thus serve as negative control. SBD1-2^C^ and SBD1-Δ#-SBD2^C^ mutants labelled with Cy3B- and ATTO647N maleimide did not show any change in stoichiometry in any case (Figure S11). We emphasize that the shown experiments serve as a proof-of-concept that PIFE-FRET is applicable to protein-protein interactions, yet no further detailed interpretations or mechanistic conclusions could be drawn from the data.

## DISCUSSION

We have here presented a detailed study of the biochemical, biophysical and structural properties of tandem substrate-binding domains of the amino acid importer GlnPQ from *L. Lactis*. We determined the crystal structure of the tandem SBDs without ligand. The packing of protein molecules in the crystal may not reflect the domain orientation of SBD1 and SBD2 in solution, but at the same time, the structures of individual SBD1 and SBD2 were in excellent agreement to the published structures of single SBDs [8]. In each tandem of the asymmetric unit (chain A/B), we observed different interactions between the domains and/or the domains and linker (Figure). Although the two observed distinct packing modes might be crystallization artefacts, it nevertheless pinpoints to an inherent mobility provided by the linker, which allows different interactions between residues of both domains (Figure). Furthermore, chain conformation A proved to be a useful tool for prediction of dye-protein interactions for assay design (Figure 7/8).

**Figure 7.**
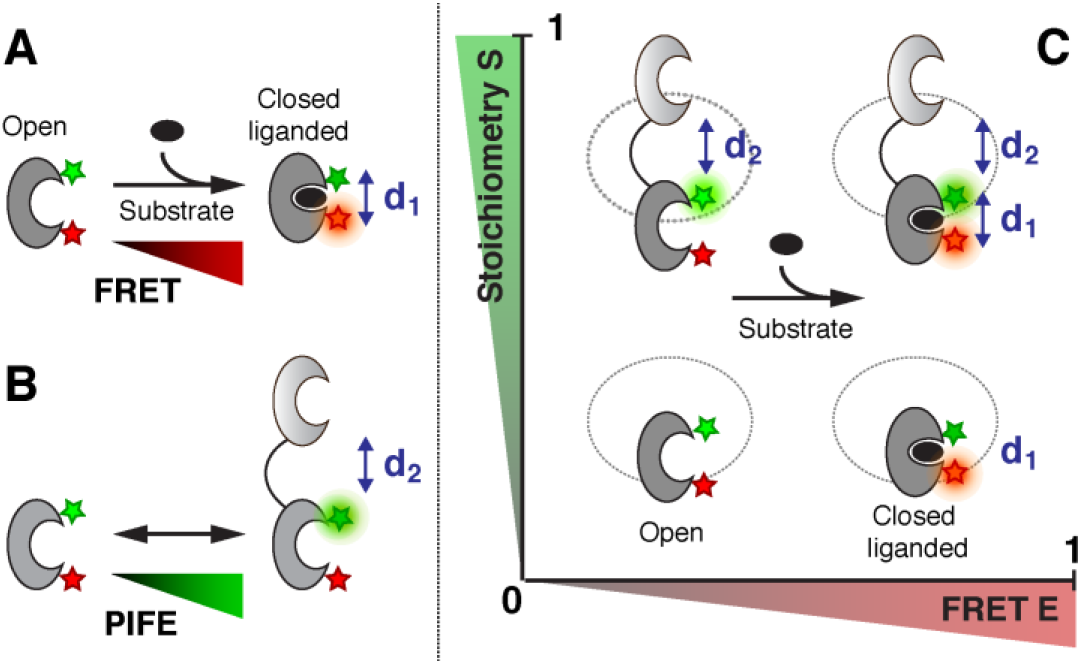
Working principle of PIFE-FRET in tandem-linked SBD1-2. **(A)** FRET between a donor and acceptor molecule reports on distance changes d1 within one of the SBDs. **(B)** PIFE can additionally report on the distance between the linked SBDs. **(C)** Readout of PIFE and FRET via ALEX spectroscopy: in a schematic E-S-histogram, conformational changes d1 within the SBD are monitored via the FRET efficiency E, whereas the distance d2 between the two SBDs is determined by changes in stoichiometry S.

**Figure 8.**
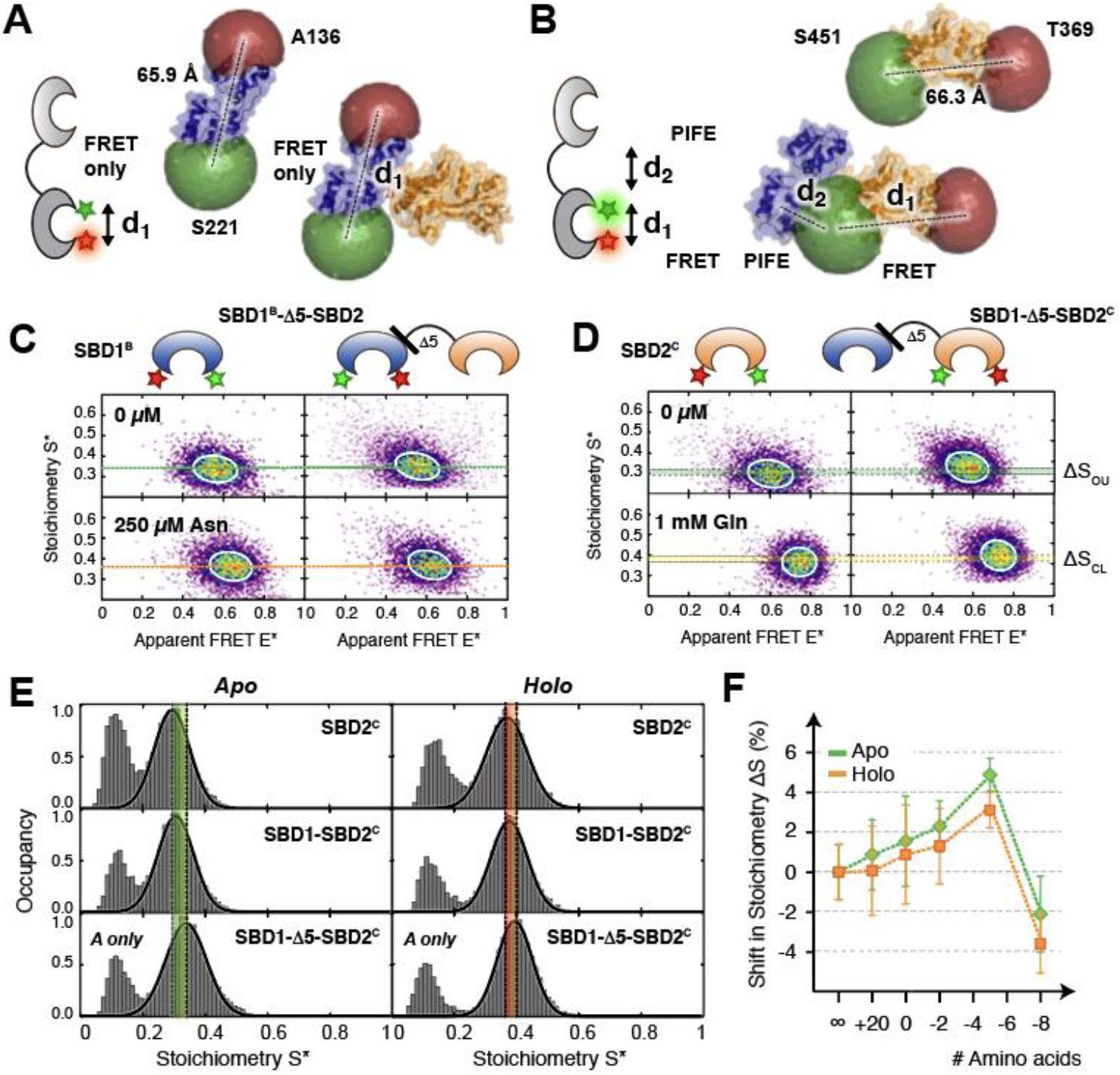
Tandem SBD1-2 investigated by PIFE-FRET. **(A-B)** Structure of SBD1-2, including labelling positions and accessible volumes of the dyes. **(A)** AV simulations predict no contact between the donor fluorophore on SBD1 – neither at A136 nor S221 – and SBD2. Note that this does not consider the case where the linker allow flexible rotation of both domains. **(B)** In SBD2, however, the donor fluorophore at S451 is affected by the presence of SBD1, hence leading to PIFE of the fluorophore. **(C)** ALEX spectroscopy on SBD1^B^ and SBD1^B^-Δ5-SBD2 labelled with Cy3/ATTO647N in the presence and absence of asparagine. No shift in stoichiometry is observed between both mutants in open/closed state. **(D)** ALEX spectroscopy on SBD2^C^ and SBD1-Δ5-SBD2^C^ labelled with Cy3/ATTO647N in the presence and absence of glutamine. A shift in stoichiometry is observed between SBD2^C^ and in SBD1-Δ5-SBD2^C^ for both, open unliganded and closed liganded state. **(E)** Stoichiometry histograms of SBD2^C^, SBD1-Δ5-SBD2^C^ and SBD1-Δ5-SBD2^C^ in the *apo* and *holo* state labelled with Cy3- and Atto647N-maleimide. The Stoichiometry value increases depending on the presence and distance to the second domain SBD1. **(F)** Shift in stoichiometry as a function of linker length. Mean values and standard deviations of the open (closed liganded) state represent the average of 2-4 independent experiments (see SI, Table S8).

**Figure 9.**
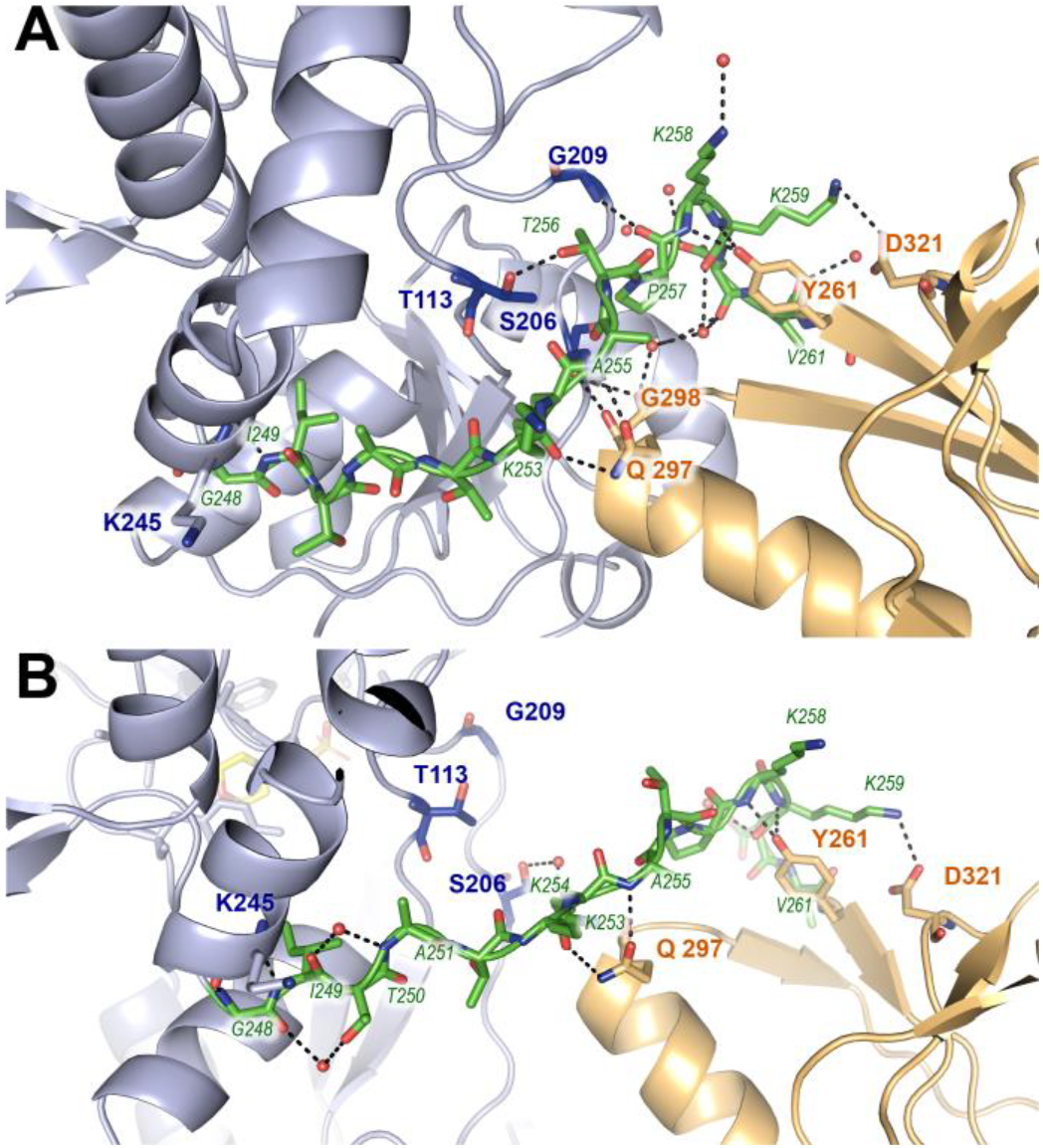
Interaction between the SBDs and their connecting linker in the crystal. Comparison between the two crystallographic monomers in the unit cell. SBD1 is depicted in blue, SBD2 in orange and linker in green**. (A)** interactions in the compact conformation (chain A) and **(B)** in looser conformation (chain B).

To examine whether the linker does indeed impart flexibility and how the tandem arrangement might impact ligand affinity, we turned to ITC and in-solution smFRET experiments. We could show that the in solution unliganded and closed-liganded states of single and tandem SBDs and SBDs were identical. In line with this observation, a combination of smFRET and ITC experiments revealed similar ligand affinities for the single SBDs in comparison to those in the tandem SBD1-2. We can thus conclude from this data that the interactions between the domains and the linker do not alter the domain structure and have little impact on the kinetics of conformational transitions since the binding constants are primarily determined by the closed-state lifetimes of the SBDs. Our experiments, however, were not able determine whether the orientation of the domains is free to change in solution or what might be the relevance of the different domain orientations seen in the crystal structure.

We further find that deletions and mutations of residues in the linker of wildtype SBD1-2 do not impact the biochemical properties or conformational states of the SBDs very much. Only in the extreme case of the short linker in SBD1-Δ5-SBD2, where 5 amino acids were removed, could we infer additional interactions between SBD1 and SBD2 that may explain the observed change in binding affinity in SBD1. For SBD1, the hinge between the domains is important for the Venus-Fly trap motion during substrate binding. The linker is directly positioned at the back of the hinge with which it can easily interact and thereby alter the binding characteristics. Similar behaviour has also been reported for other substrate-binding proteins, such as MalE[44]. In the case of SBD1-Δ5-SBD2, the shortening of the linker might alter the orientation or rotation possibilities of SBD2 relative to SBD1, which may be reflected in altered biochemical parameters, i.e., higher affinity. In line with results from ITC experiments, we further observed in our smFRET assays, that the binding affinity of glutamine to SBD2 in SBD1-Δ5-SBD2^C^ is decreased (Figure S10). This further supports the hypothesis that SBD1 and SBD2 are no longer connected in a flexible fashion and might force SBD2 to slightly open in SBD1-Δ8-SBD2^C^ seen by a shift in mean FRET efficiency (Figure S9/S11).

To further probe inter-domain interactions and to provide a proof-of-concept experiment, we implemented solution-based PIFE-FRET [29, 30] showing that it is possible to simultaneously monitor conformational states and binding affinities of one SBD while probing the proximity to the neighbouring protein domain. We have previously shown that the combination of PIFE and FRET allows probing both, short and long distances in protein-nucleic acid interactions in a single experiment. In the experiments shown here, we monitor protein-protein interactions via PIFE-FRET and provide a proof-of-concept to probe the proximity and influence of an unlabelled protein domain to a neighbouring simultaneously via smFRET.

Our data also suggest that the native coupling of both domains has no functional significance for ligand-binding. We show that the tandem SBD1-ε20-SBD2 in solution has identical conformational states and binding properties as the wildtype tandem SBDs (Figure S11). In line with this, we do not find positive or negative cooperativity in the binding of ligand to one or the other SBD. Yet, the linker length is an important factor for transport activity. For instance, extensions of the linker resulted in a reduced rate of transport, presumably by increasing the delivery time for ligand from the SBDs to the TMDs of the transporter [17]. Our work clearly shows that the ligand binding properties of each SBD are nearly unaffected by the neighbouring domain.

## METHODS AND MATERIALS

### Preparation of reagents

Unless otherwise stated, reagents of luminescent grade were used as received. Ingredients for buffers as well as chemical compounds such as 6-hydroxy-2,5,7,8-tetramethylchro-mane-2-carboxylic acid (Trolox), dithiothreitol (DTT), EDTA, bovine serum albumin (BSA), asparagine and glutamine were purchased from Sigma-Aldrich. The radio-labelled compounds [^3^ H]-asparagine and [^14^ C]-glutamine were obtained from American Radiolabeled Chemicals and PerkinEllmer, respectively. Recombinant DNA reagents and primers were purchased from Merck. As calibration samples in ALEX as well as anisotropy experiments, 45 bp-long oligonucleotides (IBA, Germany) were used as received. The single-stranded DNA was labelled with Alexa Fluor 555, Alexa Fluor 647 (Thermofisher), Cy3, Cy3B (GE Healthcare) or ATTO647N (ATTO-Tec, Germany). Complementary ssDNA strands containing an donor and acceptor fluorophore were annealed [29] as FRET standards and stored at 100 μM in 20mM Tris-HCl (pH 8.0), 500 mM NaCl, 1mM EDTA. For these calibration experiments based on dsDNA, an imaging buffer based on PBS with 2mM Trolox at pH 7.4 [45, 46] was employed.

### Nomenclature of GlnPQ derivatives

GlnPQ is composed of two subunits: GlnP and GlnQ. GlnP corresponds to the TMD that is N-terminally linked to SBD1 and fused to SBD2 [7]. GlnQ corresponds to the NBD. In this work, we investigate the single and linked substrate-binding domains SBD1 and SBD2. To investigate the conformation, binding-kinetics and cooperativity between both SBDs based on single-molecule FRET, we mutated single SBDs by Cys residues for labelling with fluorophores. We focussed on four distinct Cys backgrounds to probe the conformational states of each substrate-binding domain and to monitor the interaction between both SBDs within the tandem. Throughout the manuscript, we omit the site-specific labelling position and denote them by superscripts. To probe conformations within SBD1, we studied A: SBD1(T159C/G87C) and B: SBD1(A136C/S221C) including their tandem mutants e.g., SBD1(T159C/G87C)-2. In case of SBD2, we focus on C: SBD2(T369C/S451C) as single domain or part of the tandem. The cysteine-background of the interdomain mutant is T: SBD1(A136C)-SBD2(T369C). We shortly refer to them as SBD1^A^, SBD1^B^, SDB2^C^, SBD1^A^-2, SBD1^B^-2, SBD1-2^C^ and SBD1^T^-2^T^. Here, SBD1-2^C^ for example refers to the SBD-tandem with two cys-mutations at position 369 and 451 on SBD2.

#### Linker

In GlnP, both substrate-binding domains are tethered together by a flexible linker of 14 amino acids. To investigate its influence on substrate binding by SBD1 or SBD2 in the presence or absence of the second SBD, we created different cys-containing mutants with shortened respectively extended amino acid sequences between both SBDs (SI, Fig. S6A). These are based on the Cys derivatives SBD1^B^, i.e., SBD1(A136C/S221C) and SBD2^C^, i.e., SBD2(S369C/S451C), respectively. We denote them as SBD1^B^-Δ#-SBD2 and SBD1-Δ#-SBD2^C^, where Δ# denotes the number of deleted amino acids; εAA denotes the number of inserted amino acids (SI, Fig. S6B). A complete list with the short and full nomenclature of all designed cys-containing mutants with native and altered linker length is provided in Supplementary Figure S6C and Table S3. Mutants with shortened linker (position 248-261, Supplementary Figure S6C–D) were created by removing amino acids after position 251 of the GlnP gene sequence. Mutants with extended linker (SI, Figure S6C–D) were designed by insertion of amino acid sequence *gggsgggsgggsgggsaaql* into linker sequence between position 251 and 252. Additional point mutations such as D417F that prevent or SBD1 or SBD2 from closing and substrate binding, are added at the end of the domain in brackets. SBD1-Δ5-SBD2^C^(D417F) refers to the tandem-protein with cysteine mutations at SBD1 and a point mutation at D417 in SBD2.

### Bacterial strains, plasmids and growth conditions

The soluble substrate-binding domains were expressed in *E. coli* strain MC1061 carrying pBADnLicSBD1 and pBADnLicSBD2 and derivatives (site-directed mutants in either SBD1 or SBD2). The cells were grown in Luria-Bertani medium supplemented with 100 μg/ml of ampicillin in shake flasks. Expression was triggered at an OD_600_ of 0.5-0.6 by adding 2×10^−4^ % L-arabinose and fermentation was continued for another 2 hours. Cells were harvested by centrifugation (15 min, 6000x*g*) and washed once with 100 mM KPi (pH 7.5). After resuspension in 50 mM KPi (pH 7.5), 20 % glycerol and the addition of 0.1 mg/ml DNase, 1 mM MgCl_2_ and 1 mM phenylmethanesulfonyl fluoride (PMSF), the cells were disrupted by sonication. After the addition of 5 mM EDTA, the supernatant (after disruption of the cells) was collected after ultracentrifugation (90 min, 150,000x*g*). The cell lysate was stored in aliquots at −80 °C after flash freezing in liquid nitrogen until used for purification.

### Cloning and mutagenesis

The genes encoding the soluble SBDs were cloned into pBADnLIC [47], using ligation independent cloning, resulting in an N-terminal extension of the proteins with a 10-His-tag and a TEV protease site as described [47]. Site-directed mutagenesis was accomplished by the uracil excision-based cloning method, which employs pfuX7 polymerase [48]. Mutations were verified by sequence analysis (Eurofins Genomics, Germany).

### Purification of SBD1 and SBD2 mutants

The cell lysate was thawed and mixed with 50 mM KPi (pH 8.0), 200 mM KCl, 20 % glycerol (buffer A) plus 20 mM imidazole and incubated with Ni^2+^-Sepharose resin (GE Healthcare Buckinghamshire, United Kingdom) (5.5 ml bed volume of Ni^2+^-Sepharose was used per g of wet weight cells) for 1 h at 4 °C (under mild agitation). Next, the resin was washed with 20 column volumes of buffer A supplemented with 50 mM imidazole. The His-tagged proteins were eluted in 3 column volumes of buffer A supplemented with 500 mM imidazole. Immediately after elution and concentration determination, 5 mM EDTA was added to prevent aggregation of the proteins. The His-tag was cleaved off by His-tagged-TEV protease treatment at a ratio of 1:40 (w/w) with respect to the purified protein, and, subsequently, the protein was dialyzed against 50 mM Tris-HCl (pH 8.0), 0.5 mM EDTA plus 0.5 mM DTT overnight at 4 °C. The His-tagged TEV and residual uncut protein were removed using 0.5 ml bed volume Ni^2+^-sepharose. The flow-through of the column was concentrated (Vivaspin®, mwco 10 or 30 kDa for single SBD or tandem mutants, Sartorius; ~ 5 mg/ml), dialyzed in buffer A supplemented with 50 % glycerol, split in aliquots and stored at - 80 °C after flash freezing. Before experiments, all proteins were further purified using size-exclusion chromatography on a Superdex-200 column (GE Healthcare, Buckinghamshire, United Kingdom). Their corresponding elution profiles are shown in Figure S12. Single SBDs elute around 17.5 ml and are discernible from tandems that feature an accelerated elution around 15.5 ml. All fractions of the eluted proteins were collected (Figure S12), and re-concentrated prior to fluorescence labelling for single-molecule experiments. The column was equilibrated in 50 mM Kpi (pH 8.0), 200 mM KCl in case of subsequent ITC and DSC experiments, or 20 mM Hepes-NaOH (pH 7.5), 150 mM NaCl in case of crystallization.

### Crystallization and structure determination

SBD1-2 crystals were grown with the hanging drop vapour diffusion method at 281 K [49]. Drops were prepared by mixing the protein (concentrated to 23 mg/ml) and reservoir solution in a 1:1 v/v ratio. Crystals grew from a reservoir solution containing 125 mM MES (pH 6.0), 25% PEG200, 6.25% PEG3350 plus 50 mM NaF within 1-3 days. Data were collected at the beamline ID14-1, ESRF, Grenoble, France. The recorded data were processed using XDS [50] software package and revealed, that SBD1-2 crystals belong to space group C222_1_ with two molecules per asymmetric unit cell and 58% solvent content. The structure was solved by molecular replacement, using Phaser 2.1.4 as part of the CCP4 program suite [51]. To solve the unliganded structure for SBD1-2, the structure of the single domains were used (PDB 4KQP for SBD2 and 4LA9 for SBD1 [8]). The model building and corrections were carried out using the program COOT [52]. The models were refined using Phenix [53] with 5% of reflection randomly set aside to monitor the refinement progress. The overall quality of the model was assessed using the program MolProbity [54]. Final refinement statistics are shown in Table S1. The tandem SBD1-2 unliganded has been deposited to the PDB bank with the PDB ID 6H30.

### Isothermal Titration Calorimetry

Isothermal Titration Calorimetry (ITC) experiments were performed as described previously [8]. Briefly, the purified SBDs were dialyzed overnight against 50 mM KPi (pH 6.0), 1 mM EDTA and 1 mM NaN_3_. Isothermal titration experiments were carried out using an ITC-200 (Micro Cal™, GE Healthcare, Buckinghamshire, United Kingdom). For these experiments, the substrate was prepared in the dialysis buffer to minimize mixing effects. All experiments were carried out at 25 °C and a mixing rate of 1000 rpm. The concentration of SBD1 and SBD2 and associated tandem mutants varied between 20 and 100 μM during the experiment, depending on the expected K_D_ of the protein under investigation. For titration experiments with asparagine typically the concentration of ligand in the syringe was 8 – 10x the concentration in the cell. For glutamine titrations to SBD2, the concentration in the syringe varied between 8 and 40x protein concentration. The recorded data was approximated by a one- respectively two-site binding model [55] and fitted using the nonlinear curve-fitting tool provided by ORIGIN 8 (Origin Lab Corp. Northhampton, MA) to describe the molar enthalpy change ΔH for protein-ligand complex formation, the stoichiometry *n* and the corresponding association constant K_A_. From these, we derived the dissociation constant K_D_ as 1/K_A_, and the standard free energy change of binding ΔG = − RT ln(*K_A_*). The molar entropy change ΔS was calculated from ΔG = ΔH - TΔS. The experiments were at least repeated 3 times, if not mentioned otherwise. For analysing of the glutamine binding to SBD1-Δ5-SBD2, which features two binding sites, we first determined the parameters for the single site using the conditions of asparagine binding to SBD1. Next, the analysis of the SBD1-Δ5-SBD2 mutants for the titration with glutamine was performed. Afterwards, we fixed the parameters for SBD2 for the 2-site-fitting model in order to determine the binding of glutamine to SBD1.

### Differential Scanning Calorimetry

To determine the proteins thermal stability, differential Scanning Calorimetry (DCS) experiments were performed as described previously [8]. Briefly, the purified SBDs were dialyzed overnight against the DSC working buffer, i.e., 50 mM KPi (pH 7.0), 150 mM KCl, 1 mM EDTA and 1 mM NaN_3_. DCS experiments with working buffer solutions containing 4 μM of an SBD-mutant and 5 mM substrate were conducted on a VP-DSC Calorimeter (Micro Cal™, GE Healthcare, Buckinghamshire). The melting temperature T_m_ was determined by ORIGIN 8 (Origin Lab Corp, Northampton, MA).

### Purification of Cys-containing mutants and protein labelling

Unlabelled SBD mutants with two inserted cysteines were stored at −20 °C in 100 μl aliquots of 20-40 mg/ml in 50 mM KPi (pH 7.4), 50 mM KCl, 50 % glycerol and 1mM DTT. Stochastic labelling with maleimide derivatives of donor and acceptor fluorophores was carried out on ~ 5 nmol of protein with a ratio of protein:donor:acceptor = 1:4:5; SBD derivatives were labelled with two dye pairs: Alexa Fluor 555- and Alexa Fluor 647-maleimide (FRET assay) or Cy3(B)- and ATTO647N-maleimide (PIFE-FRET assay). Briefly, purified proteins were treated with DTT (10 mM; 30 min) to fully reduce oxidized cysteines. After diluting the protein sample to a DTT concentration of 1 mM, the reduced protein was bound to a Ni^2+^-Sepharose resin (GE Healthcare, UK) and washed with 10 column volumes of 50 mM KPi (pH 7.4), 50 mM KCl, 10% glycerol (buffer B). Simultaneously, the applied fluorophore stocks (50 nmol in powder) dissolved in 5 μl of water-free DMSO, were added at appropriate amounts to buffer B and immediately applied to the protein bound to the Ni^2+^-Sepharose resin (keeping the final DMSO concentration below 1 %). The resin was incubated overnight and kept at 4 °C (under mild agitation). After labelling, the unbound dye was removed by sequential washing with 10 column volumes of buffer B, followed by 100 column volumes of 50 mM KPI (pH 7.4), 10 mM KCl, 5 % glycerol. The protein was eluted in 0.8 ml of 50 mM KPI (pH 7.4), 50 mM KCl, 5 % glycerol, 500 mM imidazole, and applied onto a Superdex-200 column (GE Healthcare, UK) equilibrated with 50 mM KPi (pH 7.4), 200 mM KCl.

### Steady-state fluorescence anisotropy

Free fluorophore rotation and hence the correlation between FRET efficiency and distance were validated by steady-state anisotropy measurements. Fluorescence spectra and anisotropies *R* [56] were derived on a standard scanning spectrofluorometer (Jasco FP-8300; 20nm exc. and em. width; 8 sec integration time) and calculated at the emission maxima of the fluorophores (e.g., λ_em_ = 570 nm for Cy3(B), and λ_em_ = 660 nm for ATTO647N) according to the relationship *R* = (I_VV_ − GI_VH_) / (I_VV_ + 2GI_VH_). The excitation wavelengths at λ_ex_ = 532 nm resp. λ_ex_ = 640 nm were chosen according to the laser lines employed for μs ALEX spectroscopy. I_VV_ and I_VH_ describe the emission components relative to the vertical (V) or horizontal (H) orientation of the excitation and emission polarizer. The sensitivity of the spectrometer for different polarizations was corrected using horizontal excitation to obtain G = I_HV_ / I_HH_. Typical G-values for Cy3(B) and ATTO647N were 0.64 ± 0.03 and 0.45 ± 0.03. G-values for Alexa Fluor 555 and Alexa Fluor 647 were determined to be 1.8 - 1.9 [19]. We analysed the anisotropy of double-labelled protein mutants and DNA samples in a concentration range of about ~ 100 nM. The determined anisotropy values are summarized in SI, Table S4.

### Sample preparation for single-molecule experiments

μs-ALEX-experiments were carried out at 25-50 pM of double-labelled protein or DNA in buffer containing 50 mM KPi (pH 7.4), 150 mM KCl, 1 mM Trolox and 10 mM MEA. ALEX titration experiments on GlnPQ, i.e., a chosen SBD or tandem mutant in presence of varying ligand concentrations, were completed in one continuous experiment. To monitor and detect possible changes in the experimental settings, every set of ALEX experiments on SBDs was complemented by an experiment of dsDNA FRET standard [18] of 45 bp length (data not shown). The dsDNA was labelled either with Cy3(B) and ATTO647N or Alexa Fluor 555 and Alexa 647 in 18 and 23 bp distance depending on labelling scheme of the SBDs.

### Single-molecule FRET and ALEX-Spectroscopy

μs-ALEX-experiments were carried out at room temperature (22°C) on a custom-built confocal microscope [18, 29]. In brief, alternating laser excitation (ALEX) between 532 and 640 nm was employed with an alternation period of 50 μs, coupled into a confocal microscope, a 60x objective with NA = 1.35 (Olympus, UPLSAPO 60XO) focused the excitation light to a diffraction-limited spot 20 μm into the solution. The excitation intensity amounted to 60 μW at 532 nm (≈ 30 kW/cm2) and 25 μW at 640 nm (≈ 25 kW/cm2). Fluorescence emission was collected and spectrally separated onto two APDs (τ-spad, Picoquant, Germany) with appropriate filters (donor channel: HC582/75; acceptor channel: Edge Basic 647LP; AHF Analysentechnik, Germany). The signal was recorded using a custom-written LabView program.

### ALEX data extraction and analysis

After data acquisition, the recorded fluorescence emission was analysed and processed using custom-made scripts in Python. Fluorescence photons arriving at the two detection channels (donor detection channel: D_em_; acceptor detection channel: A_em_) were assigned to either donor- or acceptor-based excitation based on their photon arrival time. From this, three-photon streams were extracted from the data corresponding to donor-based donor emission F(DD), donor-based acceptor emission F(DA) and acceptor-based acceptor emission F(AA). For each molecule diffusing through the confocal volume, fluorophore stoichiometries S and apparent FRET efficiencies E* were calculated for each fluorescent burst above a certain threshold yielding a two-dimensional histogram [36, 37]. Uncorrected FRET efficiency E* monitors the proximity between the two fluorophores and is calculated according to:

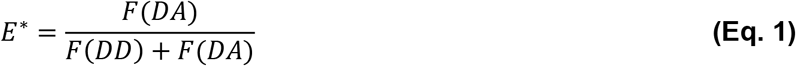

S is defined as the ratio between the overall green fluorescence intensity over the total green and red fluorescence intensity and describes the ratio of donor-to-acceptor fluorophores in the sample:

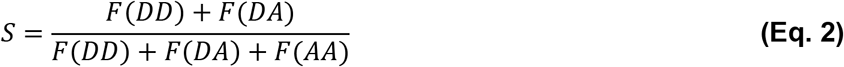

Using published procedures to identify bursts corresponding to single molecules [57], we obtained bursts characterized by three parameters (M, T, and L). A fluorescent signal is considered a burst provided it meets the following criteria: a total of L photons, having M neighbouring photons within a time interval of T microseconds. For all data presented in this study, an all photon burst search [57, 58] using parameters M = 15, T = 500 μs and L = 25 was applied; additional thresholding removed spurious changes in fluorescence intensity and selected for intense single-molecule bursts (all channels > 150 photons). After binning the detected bursts into a 2D E*/S histogram, sub-populations were separated according to their S-values. E*- and S-distributions were fitted using a 2D Gaussian function, yielding the mean values μ_*i*_of the distribution and an associated standard deviation.

### Population assignment

To correct individual populations, i.e., *apo*-protein state and closed liganded within one 2D ALEX histogram, every burst need to be assigned to a particular population. This can be achieved via cluster analysis methods or probability distribution analysis [59]. In our implementation, every population in the uncorrected 2D histogram is first fitted with a covariant bivariate Gaussian function

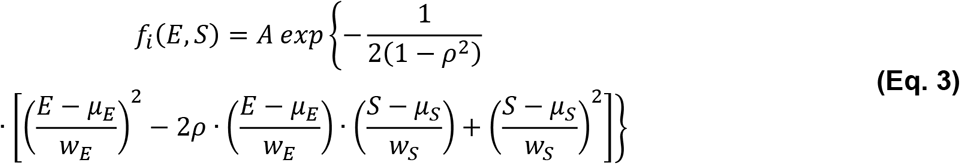

where the population is described by an amplitude *A*, its mean values *μ_i_* and standard deviations *w_i_* in FRET E and Stoichiometry S. ρ denotes the correlation matrix between E and S. We express the probability *p* that a given burst in the 2D histogram belongs to a population *i* by

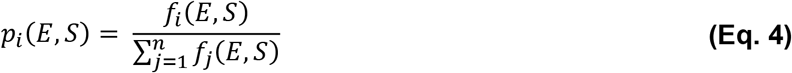

### Titration experiments

To investigate the binding affinity of the labelled SBDs, and hence the transition between open unliganded and closed liganded conformation, titrations in ALEX experiments in presence of high-affinity ligands were carried out. The two-dimensional E- and S-distributions were fitted using 2D Gaussian functions, yielding the mean values *μ_i_* of the distribution and an associated standard deviation *w_i_*. At first, the histograms of *apo*-protein and protein at fully saturating substrate concentration were investigated. Their projections in E represent the FRET distributions of the open unliganded and closed liganded state respectively. Subsequently, these two distributions were employed to fit the titration data at intermediate substrate concentration via a Hill model with fixed V_max_ value using ORIGIN 8 (Origin Lab Corp, Northampton, MA). The fractional occupancy of the high FRET Gaussian as a function of substrate concentration was fitted afterwards with a one-side-binding model, which allowed calculation of B_max_ (maximal fraction of closed state) and K_D_ (dissociation constant).

### PIFE data extraction and analysis

To monitor the presence of the second SBD and the intra-domain distance within the tandem by protein-induced fluorescence enhancement, ALEX experiments in presence of high-affinity ligands were carried out, i.e., ALEX spectroscopy on SBDs labelled with Cy3/ATTO647N-maleimide were carried without in absence and presence of a saturating ligand. The two-dimensional E*- and S*-distributions were fitted using 2D Gaussian functions (equation 3), yielding the mean values *μ_i_* of the distribution and an associated standard deviation *w_i_*. The shift in brightness of Cy3 in presence of the second SBD is seen as a shift in stoichiometry between the single SBD and as part of a tandem. Therefore, at first, the histogram of the single SDB as *apo*-protein and fully saturating substrate concentration were investigated. Their stoichiometry values are taken as reference. We report the change in stoichiometry ΔS* as a function of linker length, and hence the distance between both substrate-binding domains.

### Burst Variance analysis

To reveal any static and/or dynamic heterogeneity in single-molecule ALEX data, we employed burst variance analysis (BVA; [39]. Here, we compare the expected shot-noise limited standard deviation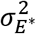 for a given mean FRET efficiency E* against the actual standard deviation for individual molecules. The expected standard deviation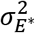 due to shot noise depends only on photon statistics and reads as:

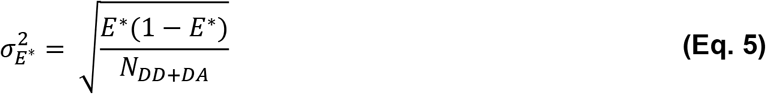

where *N_DD+DA_* is the average number of photons per burst emitted by the double-labelled molecule after green excitation. Similar burst selection criteria as described above where used: all channels >250 photons and only burst within the stoichiometry range from 0.3-0.6 were used.

### Structural modelling and accessible volume calculation

To compare distances within the obtained crystal structure with results determined by FRET, we carried out structural modelling and accessible volume calculations. We visualize the individual and linked substrate-binding domains, as well as the position at which both fluorophores are stochastically attached, based on four different crystal structures (SBD1 in the *apo*-state (PDB [8] 4LA9) and in presence of asparagine (PDB 6FXG), SBD2 in the *apo*-state (PDB[8] 4KR5) and presence of glutamine (PDB [8] 4KQP)) in comparison to the published structure of the linked substrate-binding domains. We loaded the respective pdb files in PyMOL [60] and removed co-crystallized items, like ligands and proteins. Next, we determine the ID of each CB atom to which the fluorophores, i.e., Cy3(B) resp. Alexa Fluor 555 and ATTO647N resp. Alexa Fluor 647 are attached via cysteine-maleimide click-chemistry. With this knowledge, we determined the accessible volumes (AV) and expected distances between the dyes [43] on the protein complex in the unliganded and unliganded case. The dyes were attached to the C_β_ atom of the corresponding amino acids and simulated as C2 maleimide derivative with parameters as specified in the FPS software manual [43]. Afterwards, we use PyMOL to compare and display the determined AVs and distance within the crystal structures.

## APPENDIX A SUPPLEMENTARY INFORMATION

Supplementary Information to this article can be found in the online version of this publication.

## ACKNOWLEDGEMENTS

The authors are grateful to the beam line personnel at 14-1, ID 23-1 (ESRF, Grenoble), and X06SA (SLS, Villigen) for technical assistance. We thank A.M. van Oijen for generous support of this study and R. Vietrov for fruitful discussions and advise.

## AUTHORS CONTRIBUTIONS

EP, GKS, BP and TC designed the study. BP and TC supervised the project. GKS, GG and DAG designed, overexpressed, and purified proteins. EP, GKS, NZ, AWJ and AG performed experiments and analysed data. EP, GKS, BP and TC interpreted the data and wrote the manuscript. All authors contributed to and approved the final version of the manuscript.

## Supplementary Information

### Section S1. Supplemental Figures

**Supplementary Figure S1.**
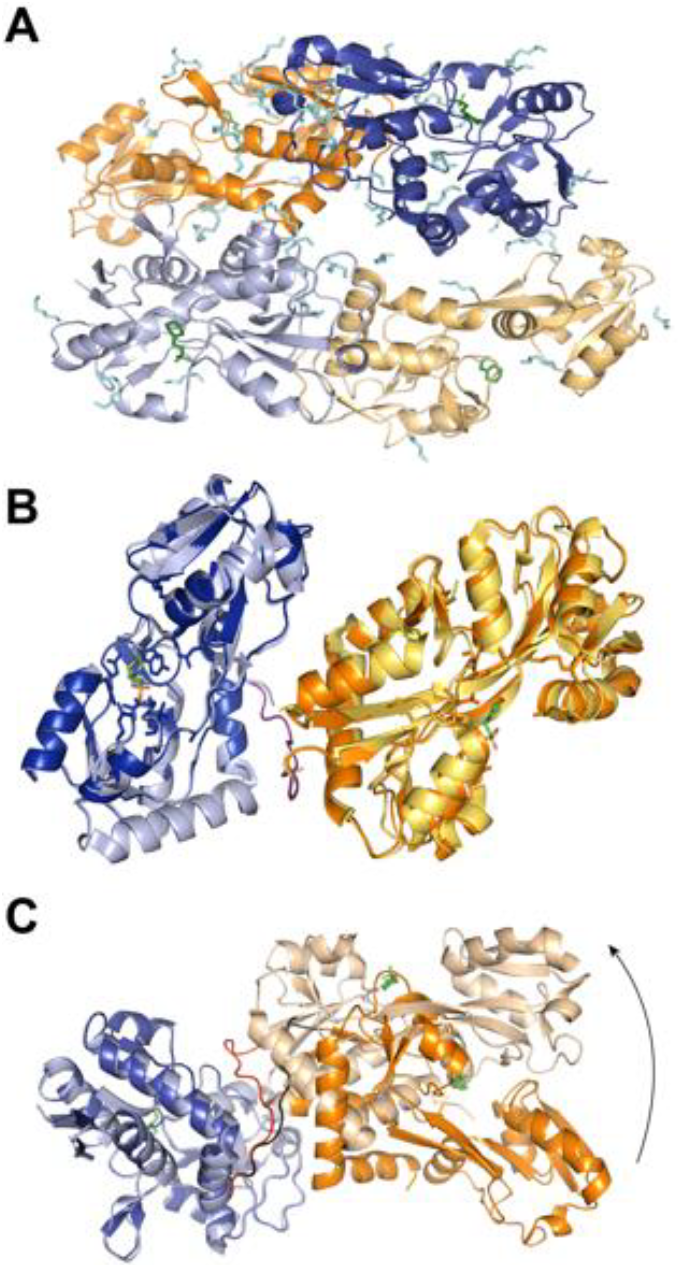
Crystal structure of unliganded SBD1-2 tandem. **(A)** Top view of the crystallographic dimer of SBD1-2 tandems. Solvent molecules are depicted in cyan. Co-crystallized MES molecules are depicted in green. SBD1 is represented in blue, SBD2 in orange. The linker (red) is close to the hinge region of SBD1. **(B)** Superposition of SBD1-2 with unliganded SBD1 (light blue; PDB ID 4KPT) and SBD2 (gold; PDB ID 4KR5). Active site residues are shown as sticks. **(C)** Superposition of both tandem proteins that are co-crystallized within C222_1_ configuration.

**Supplementary Figure S2.**
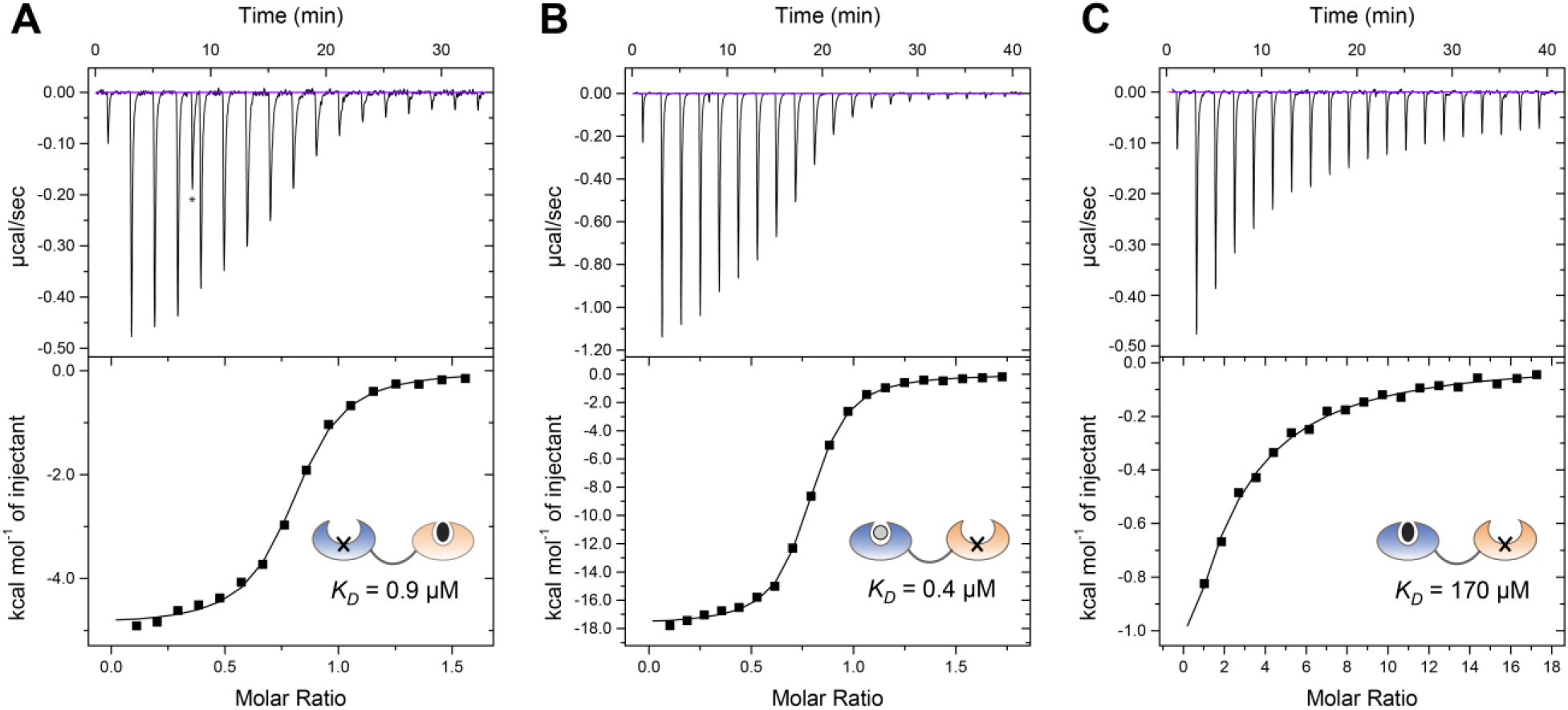
ITC data of tandem mutants with mutations in the active site. **(A)** SBD1(E184W)-2 has a K_D_ of 0.88 μM for glutamine. SBD1-2(D417F) shows a K_D_ of 0.39 μM in case of asparagine **(B)** and 170 μM in presence of glutamine **(C)**. Spikes due to leakage of the syringe (marked by asterisk) have been excluded from the analysis.

**Supplementary Figure S3.**
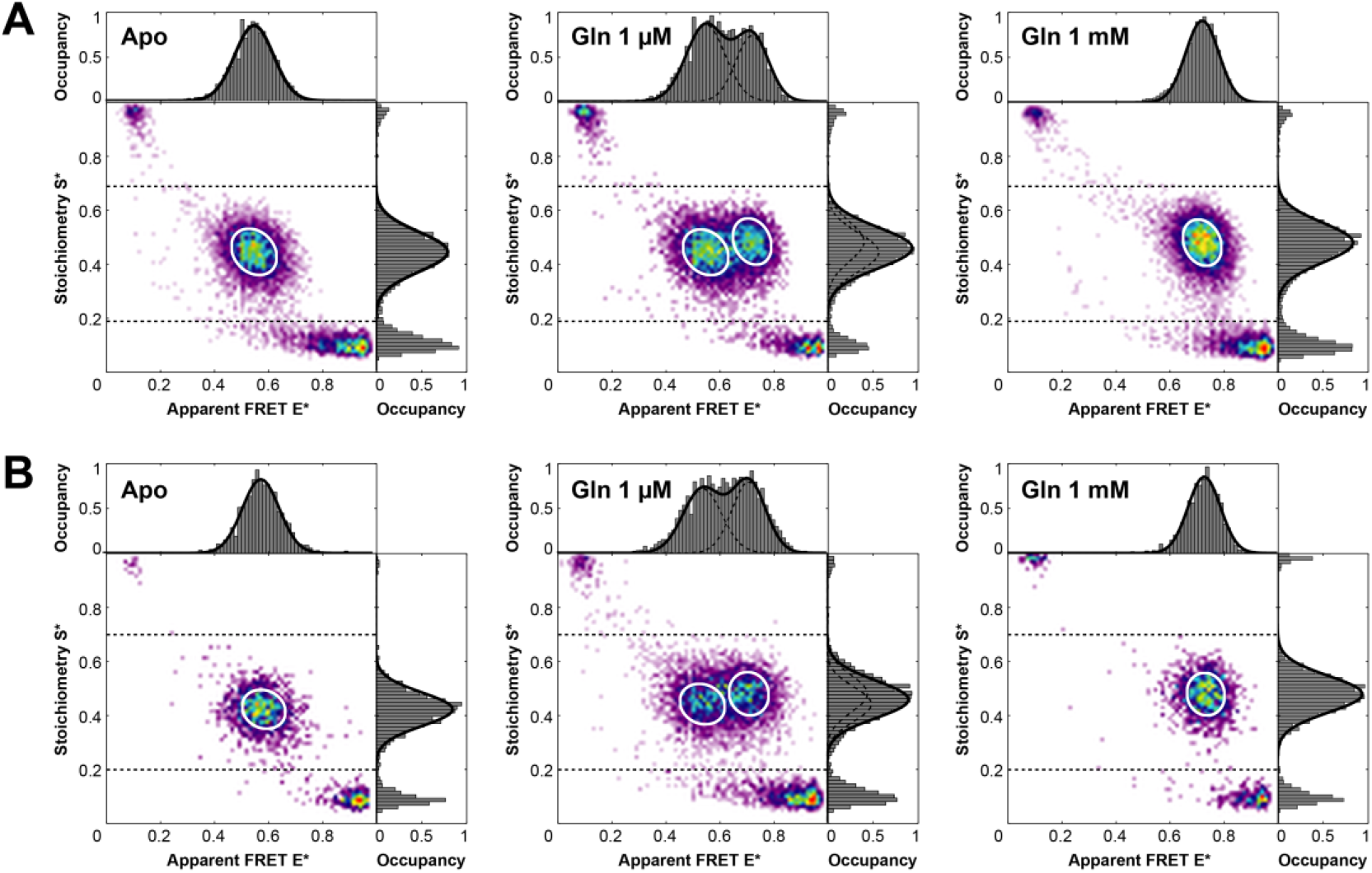
μs-ALEX Spectroscopy on isolated and fused SBD2^C^ in presence of Glutamine. ALEX-based ES-histograms of **(A)** isolated SBD2^c^ and **(B)** SBD2^C^-Tandem, labelled with Alexa Fluor 555- and Alexa Fluor 647-maleimide. Double-labelled protein species are identified in a stoichiometry range of S = 0.2-0.7. Both mutants show a low FRET value of 0.58 in the open, unliganded state and a high FRET value of 0.74 under saturating concentrations of glutamine. At substrate concentrations around the K_D_ both states are equally populated in both mutants. SBD2^C^-Tandem shows a lower occupancy of the closed-liganded state due to the presence of SBD1.

**Supplementary Figure S4.**
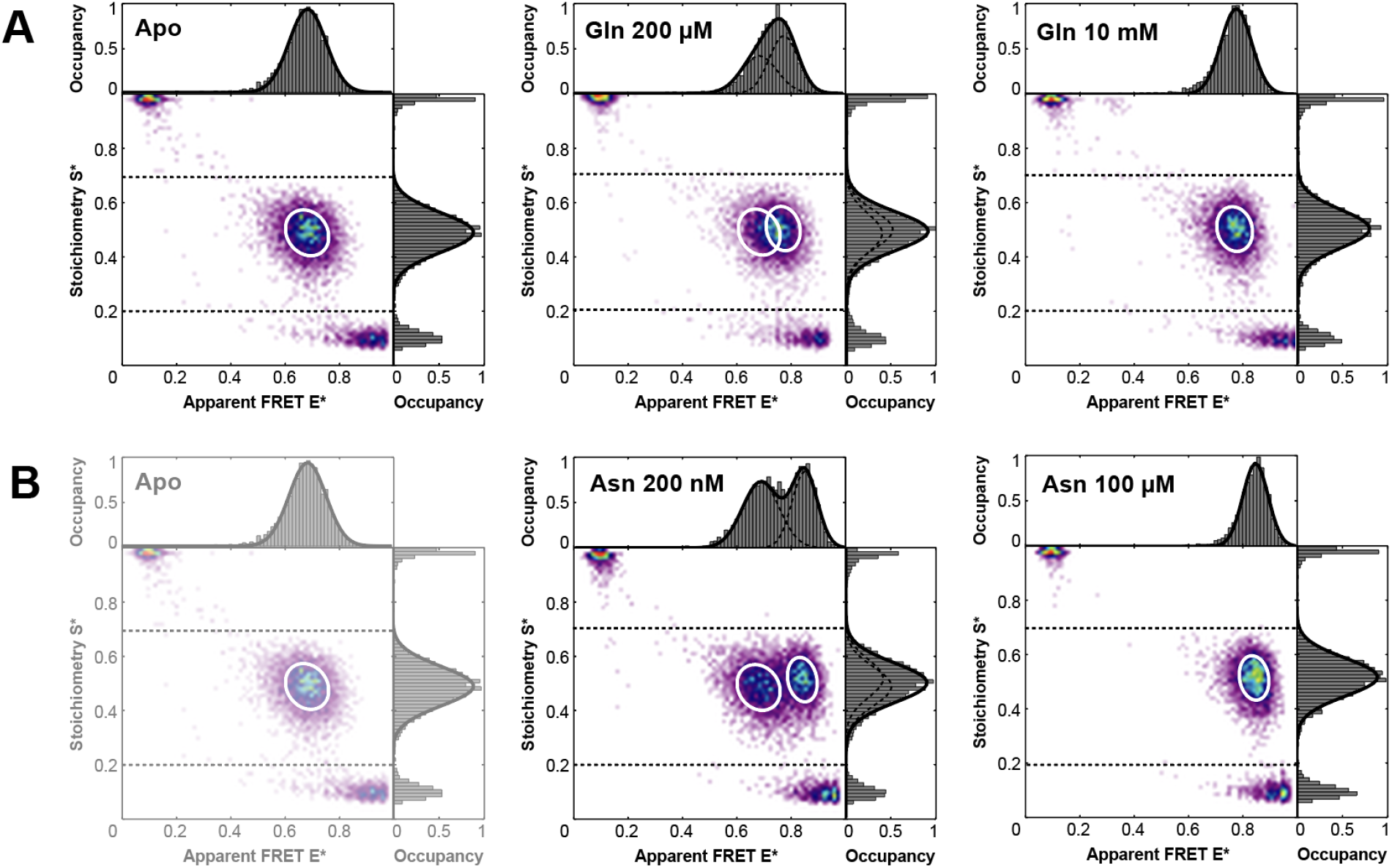
μs-ALEX Spectroscopy on isolated SBD1^A^ in presence of Glutamine, and Asparagine. ALEX-based ES-histograms of SBD1^**A**^, labelled with Alexa Fluor 555- and Alexa Fluor 647-maleimide in presence of **(A)** glutamine and **(B)** asparagine. Double-labelled protein species are identified in a stoichiometry range of S = 0.2-0.7. SBD1 shows a low FRET value of ~ 0.65 in the open, unliganded state. **(A)** For saturating concentrations of glutamine, an intermediate FRET value of ~ 0.74 is observed. At substrate concentrations around the K_D_ both states are superimposed. **(B)** For saturating concentrations of asparagine, a high FRET value of ~ 0.82 is observed. At substrate concentrations around the K_D,_ two states are equally populated.

**Supplementary Figure S5.**
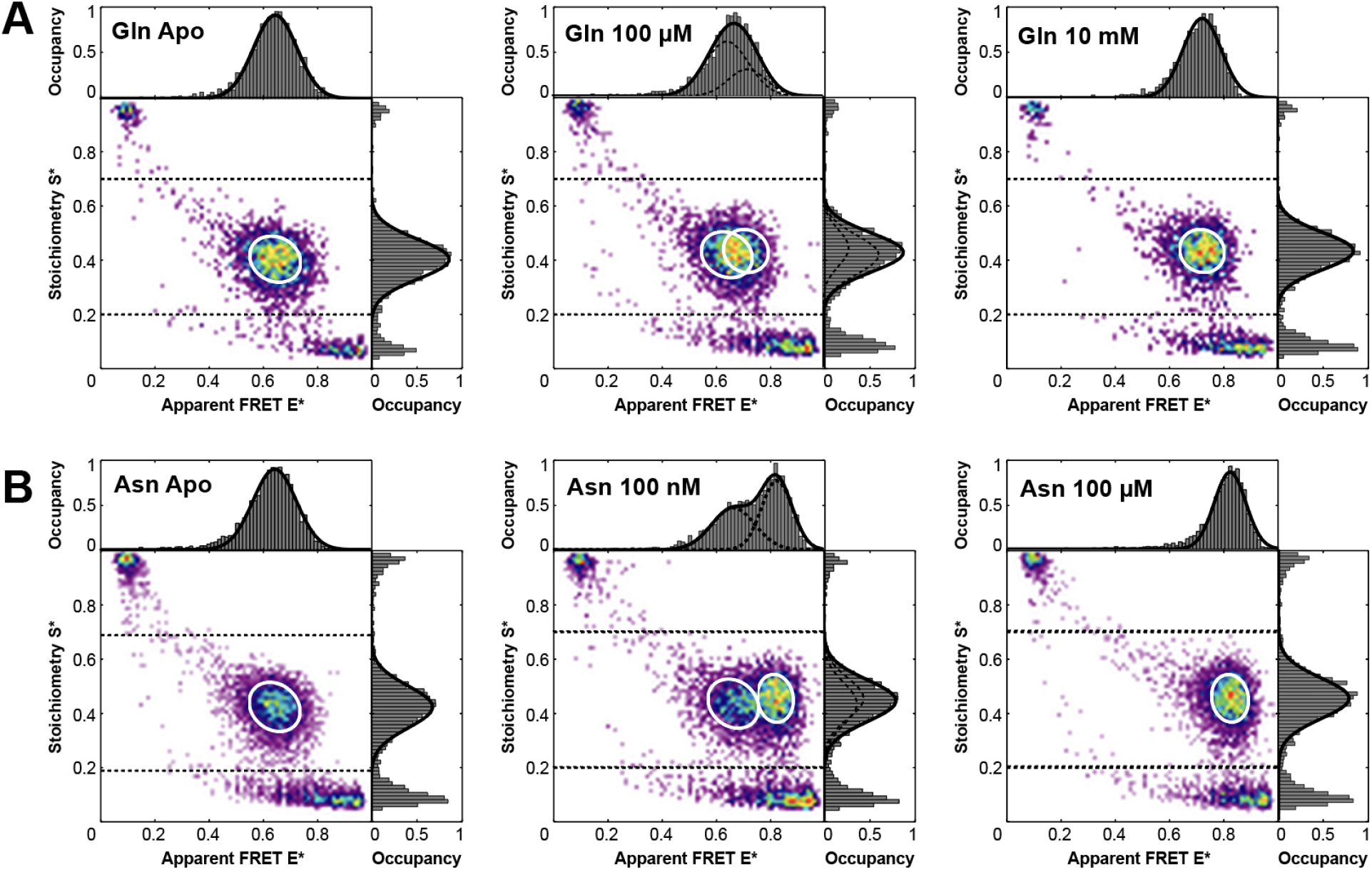
μs-ALEX Spectroscopy on isolated SBD1^A^-Tandem in presence of Glutamine and Asparagine. ALEX-based ES-histograms of SBD1, labelled with Alexa Fluor 555- and Alexa Fluor 647-maleimide in presence of **(A)** glutamine and **(B)** asparagine. Double-labelled protein species are identified in a stoichiometry range of S = 0.2-0.7. SBD1^A^-Tandem shows a low FRET value of 0.65 in the open, unliganded state. **(A)** For saturating concentrations of glutamine, an intermediate FRET value of 0.74 is observed. At substrate concentrations around the K_D_ both states are superimposed. **(B)** For saturating concentrations of asparagine, a high FRET value of 0.82 is observed. At substrate concentrations around the K_D,_ two states are equally populated.

**Supplementary Figure S6.**
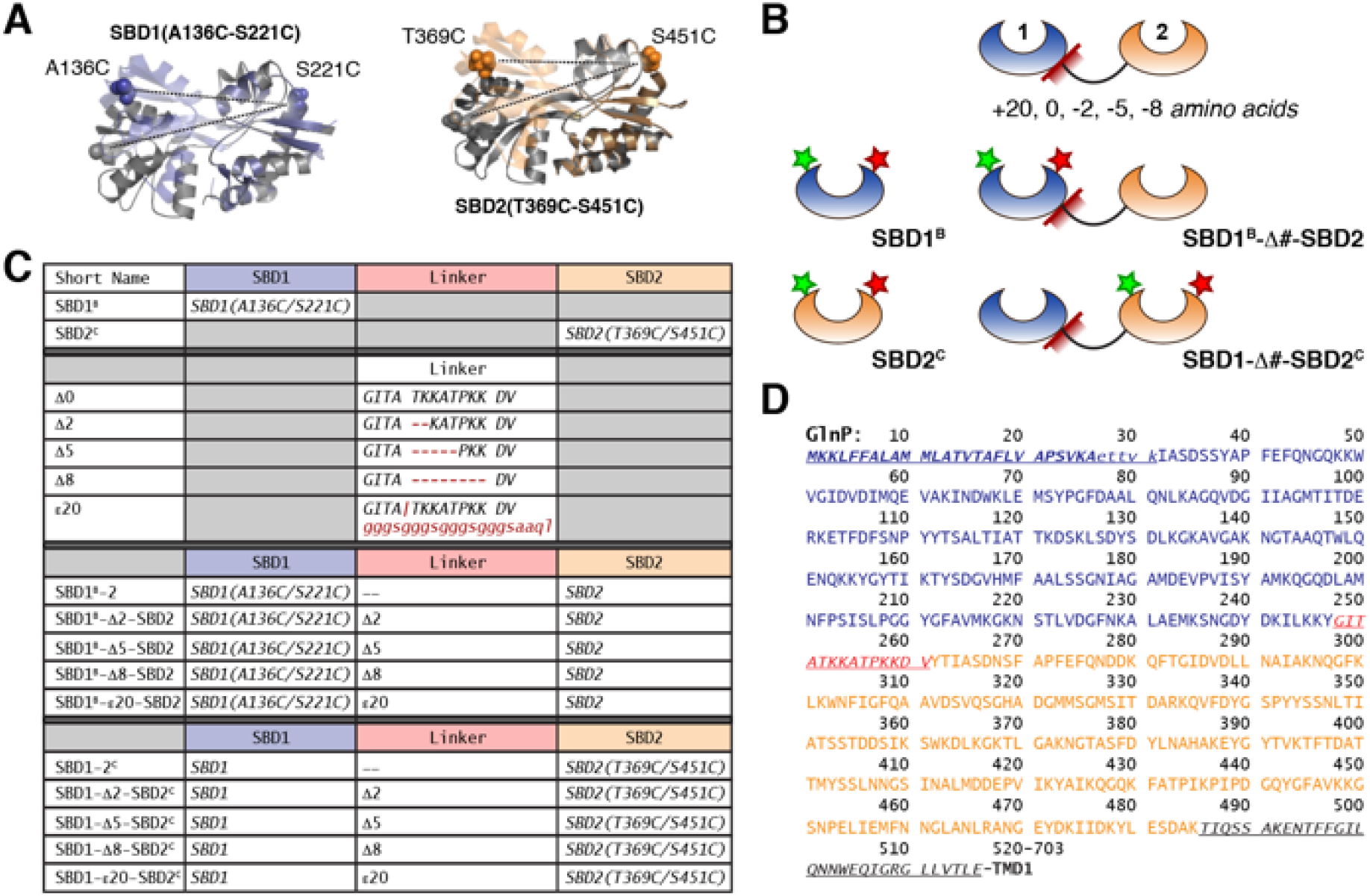
Structure and sequence of tandem mutants with linker mutations. **(A)** Crystal structures of SBD1 and SBD2 (PDB 4KR5; 4KQP; 4LA9; 6FXG). Spheres indicate the cysteine-mutations introduced for linker studies based on combined single-molecule FRET and PIFE. The open unliganded state is depicted in grey, the closed liganded state in a light colour. **(B)** Schematics of tandem mutants. **(C)** Summary of designed Cys-containing mutants with altered linker length between both substrate-binding domains. **(D)** The amino acid sequence of GlnP [1] encodes the TMD, SBD2 (orange), the linker (red) and SBD1 (blue).

**Supplementary Figure S7.**
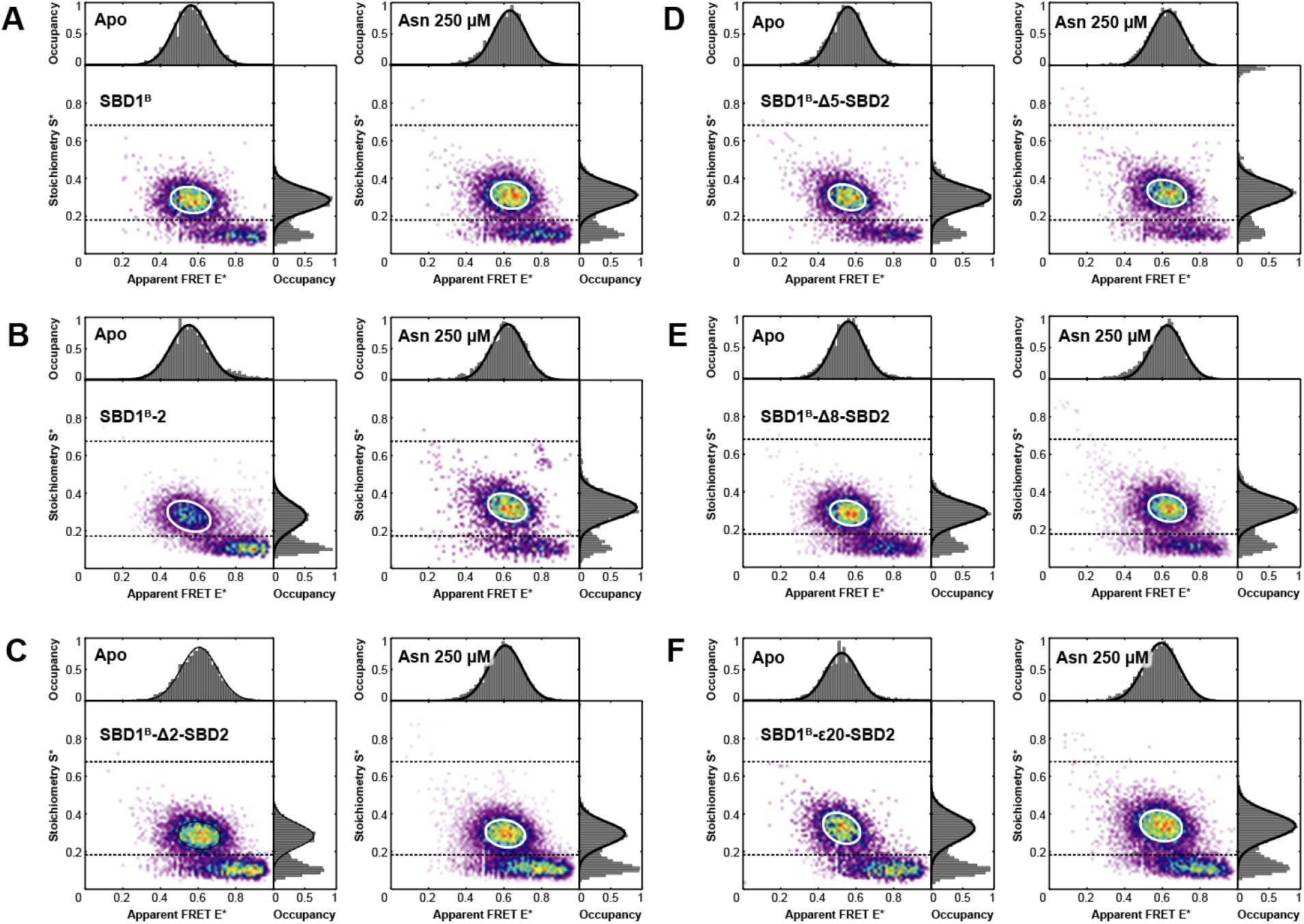
μs-ALEX Spectroscopy on SBD1^B^-Δ#-SBD2 mutants labelled with Cy3/ATTO647N maleimide. ES-histograms of proteins in the *apo-*form and in presence of 250 μM asparagine: **(A)** isolated SBD1^**B**^, **(B)** SBD1^**B**^-Tandem, **(C-F)** SBD1^**B**^-Tandem with altered linker between both domains shown for a **(C-E)** deletion of 2,5 and 8 amino acids as well as an **(F)** extension of 20 amino acids. Double-labelled protein species are identified in a stoichiometry range of S = 0.17-0.67. SBD1^**B**^-associated linker mutants show an intermediate FRET value of 0.55 in the *apo-*state. For saturating concentrations of asparagine, a FRET value of ~ 0.62 is observed. Except for SBD1^B^-Δ2-SBD2, the overall FRET shifts amount to ~ 0.06.

**Supplementary Figure S8.**
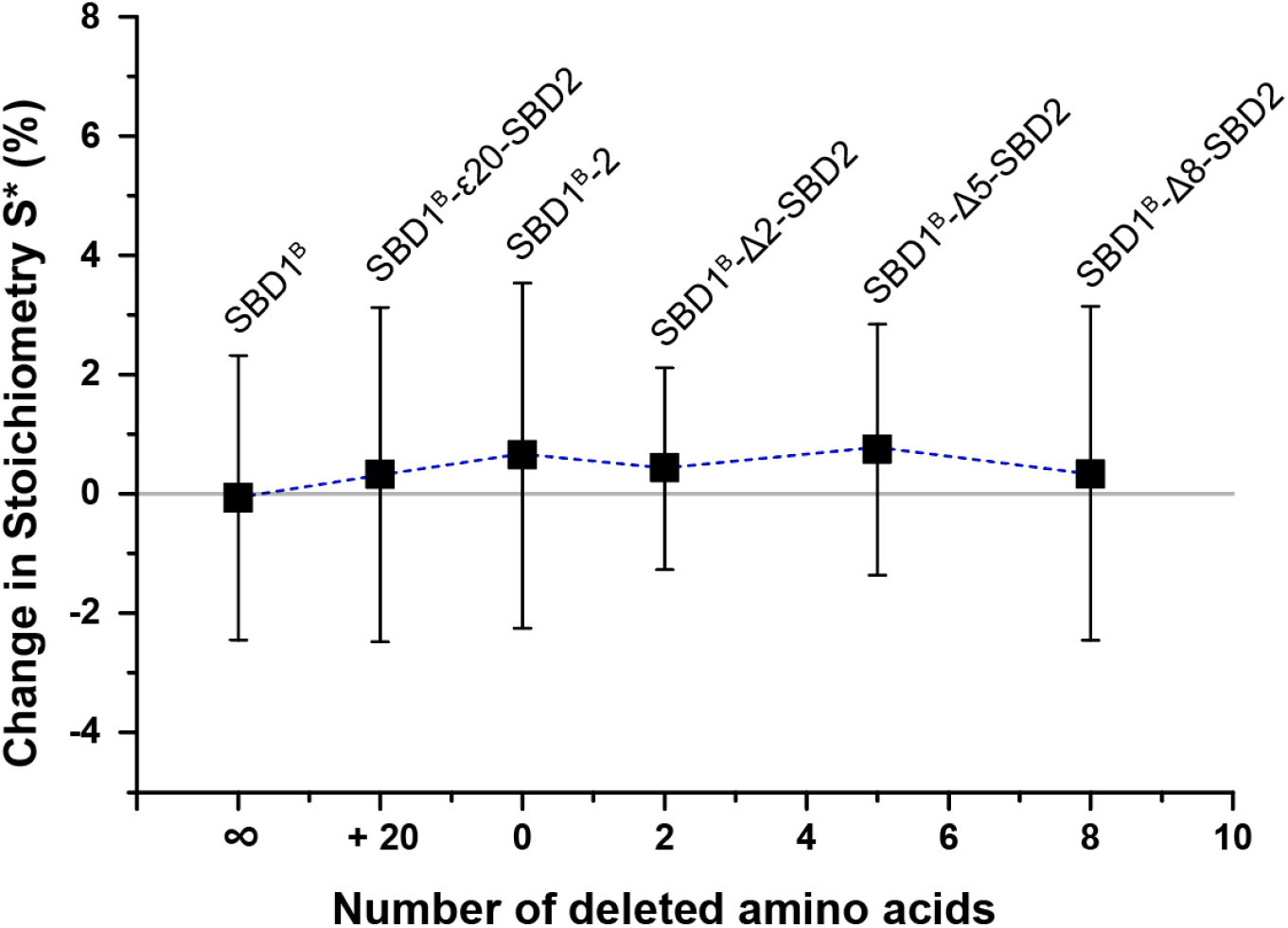
Missing enhancement due to PIFE in SBD1^B^ and SBD1^B^-tandem mutants. Shift in stoichiometry as a function of inserted or delete amino acids compared to isolated SBD1 (infinitesimal distance between both SBDs within the tandem). The stoichiometry of mutants with cysteine residues at A136 and S221 stochastically labelled with Cy3 and ATTO647N-maleimide stays constant within error bars. The attached, unlabelled domain SBD2 does not lead to a substantial spatial restriction, hence steric hindrance of the Cy3 fluorophore, which would lead to PIFE. Data represent the mean value and standard deviation of n > 3 replicas.

**Supplementary Figure S9.**
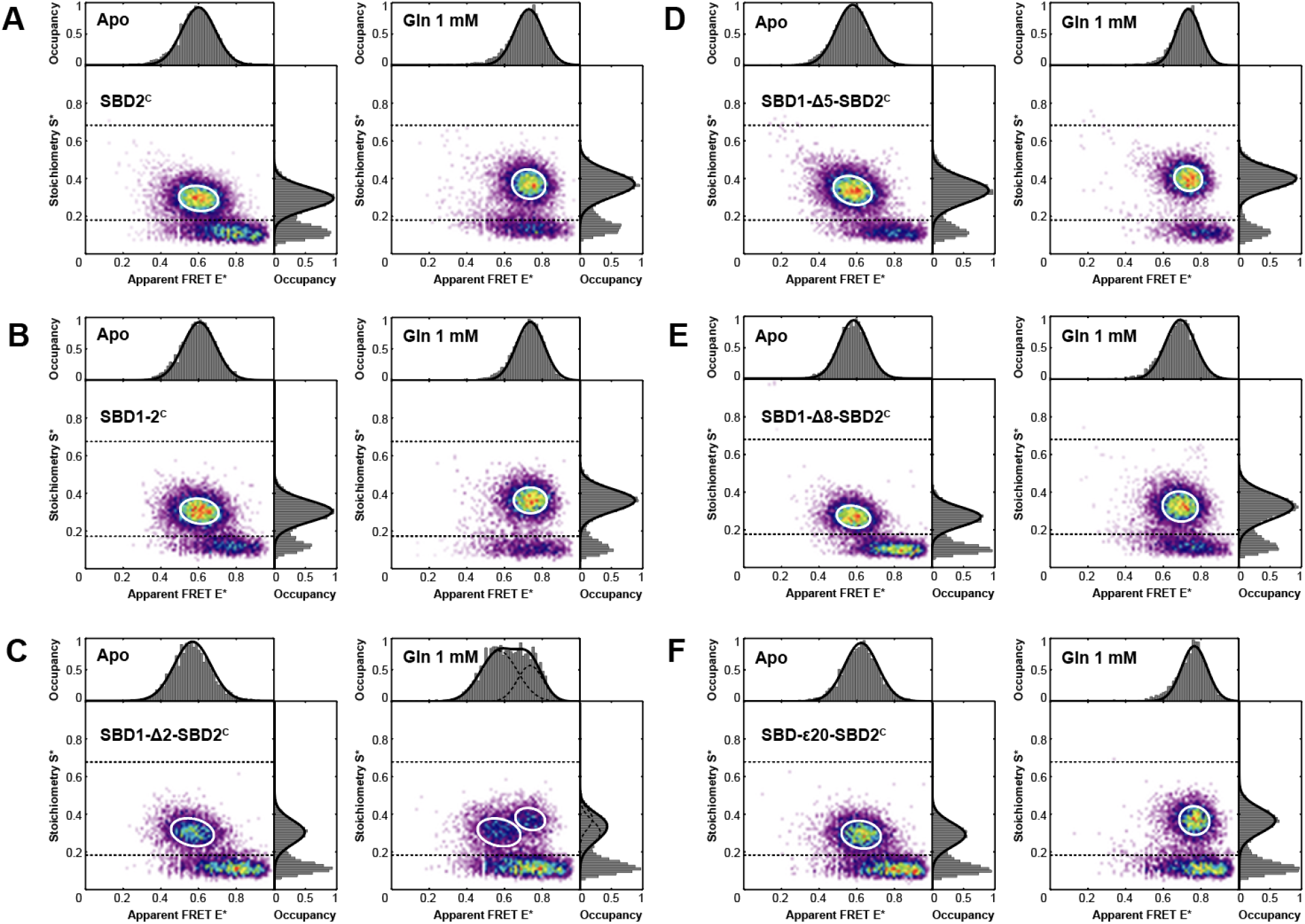
μs-ALEX Spectroscopy on SBD1-Δ#-SBD2^C^ mutants labelled with Cy3/ATTO647N maleimide. ES-histograms of proteins in the *apo-*form and in presence of 1 mM glutamine: **(A)** isolated SBD2^C^, **(B)** SBD2^C^-Tandem, **(C-F)** SBD2^C^-Tandem with altered linker between both domains shown for a **(C-E)** deletion of 2,5 and 8 amino acids as well as an **(F)** extension of 20 amino acids. Double-labelled protein species are identified in a stoichiometry range of S = 0.17-0.67. SBD2-associated linker mutants show an intermediate FRET value of ~ 0.6 in the *apo*-state. For saturating concentrations of glutamine, a FRET value of ~ 0.72 is observed. Except for SBD1-Δ5-SBD2^C^ and SBD1-Δ8-SBD2^C^, the overall FRET shift amounts to ~ 0.13.

**Supplementary Figure S10.**
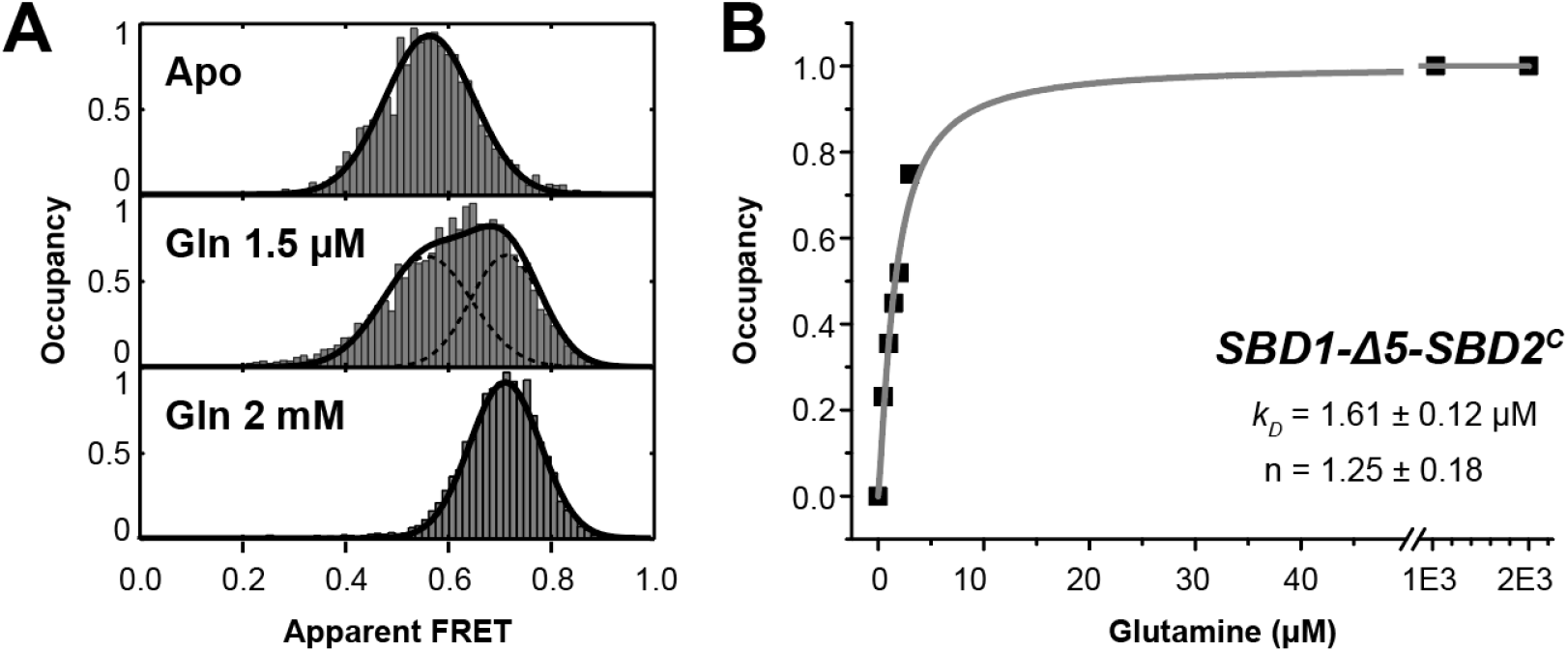
Binding affinities of the SBD2 domain as part of a tandem with shortened linker by 5 amino acids. **(A)** μs-ALEX-derived FRET histogram of SBD1-Δ5-SBD2^C^ labelled with Cy3- and ATTO647N maleimide in the *apo-*state, at concentration around the K_D_ and at saturating concentration of glutamine. **(B)** Titration curve of SBD1-Δ5-SBD2^C^ with glutamine.

**Supplementary Figure S11.**
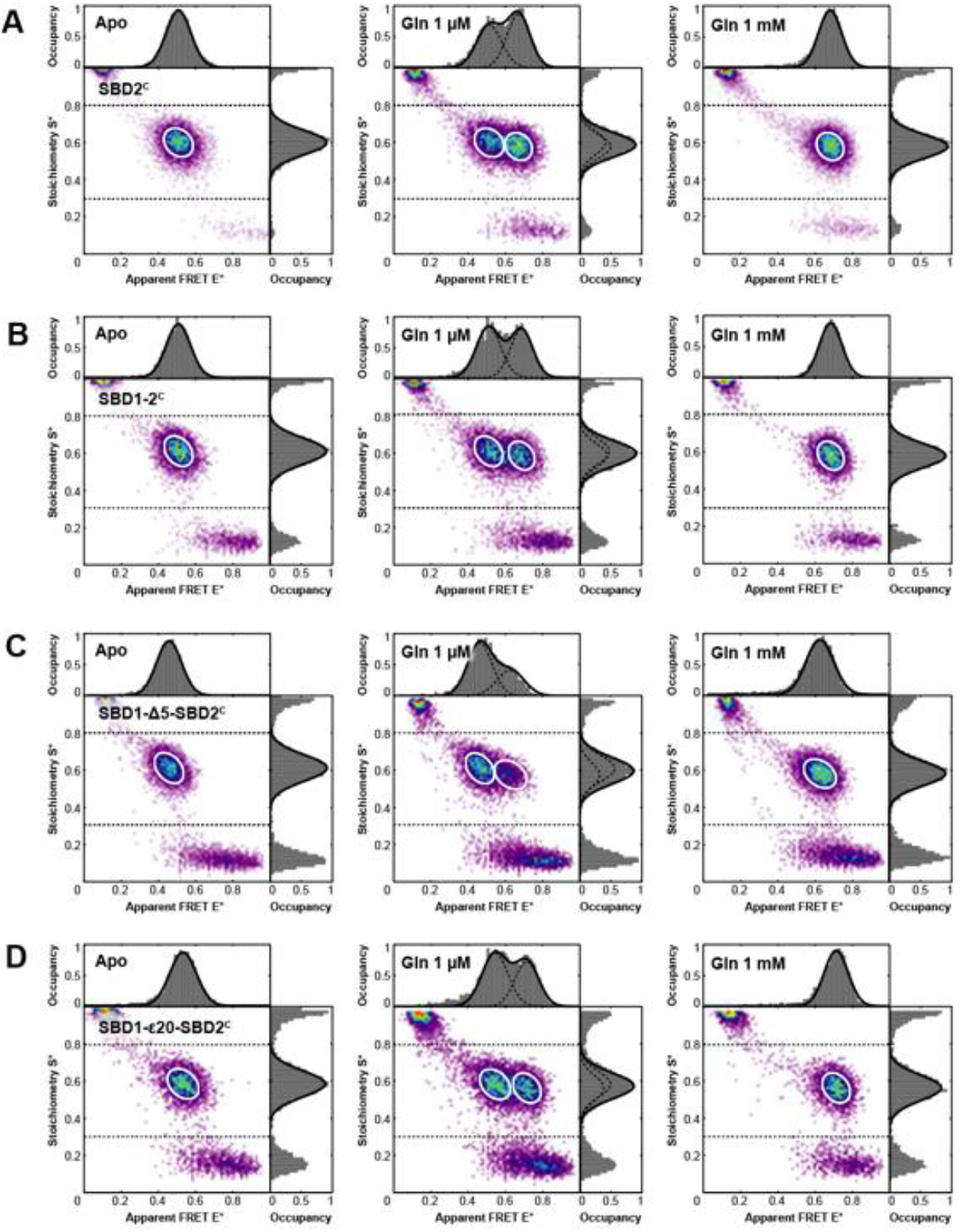
μs-ALEX Spectroscopy on SBD1-Δ#-SBD2^C^ mutants labelled with Cy3B/ATTO647N maleimide. ES-histograms of proteins in the *apo-*form, around the K_D_ and in presence of 1 mM glutamine. **(A)** SBD2, **(B)** SBD2-Tandem, SBD2-Tandem with altered linker between both domains are shown for a **(C)** deletion of 5 amino acids and **(D)** extension of 20 amino acids. Double-labelled species are identified in a stoichiometry range of S = 0.3-0.8. SBD2 associated mutants show an intermediate FRET value of 0.55 in the *apo-*state. For saturating concentrations of glutamine, a high FRET value of ~ 0.74 is observed. The overall FRET shifts amount to ~ 0.17.

**Supplementary Figure S12.**
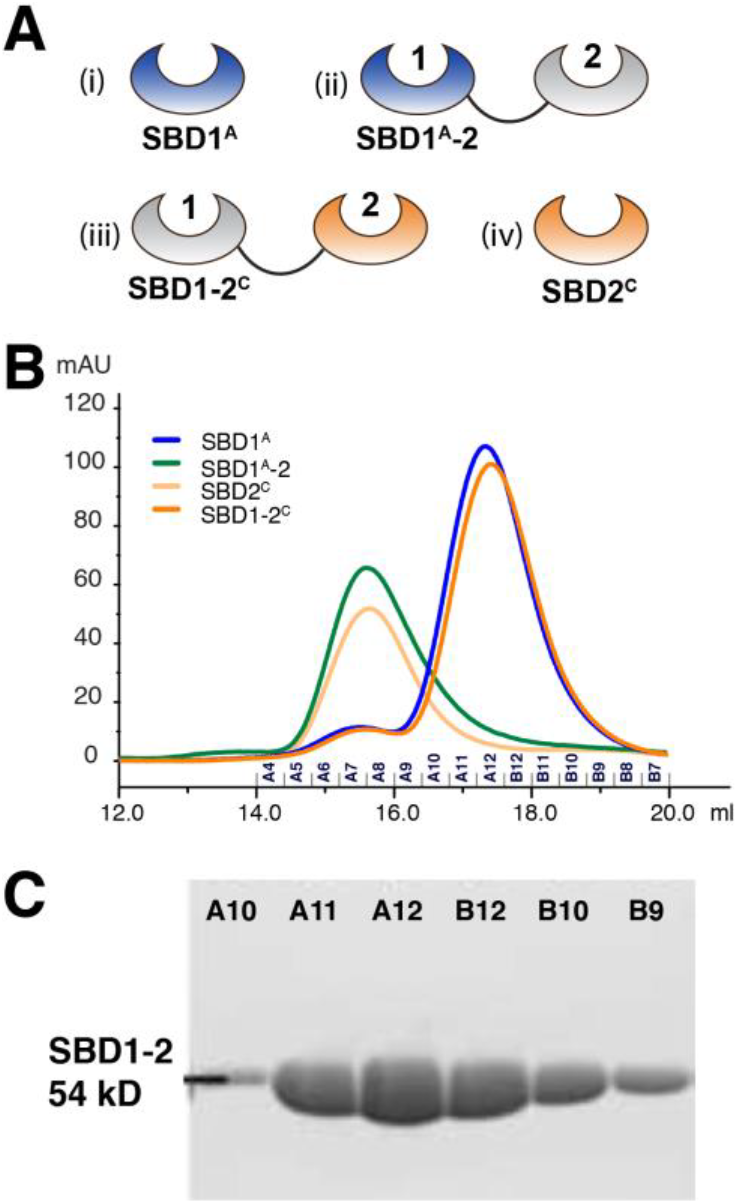
Biochemical characterization of FRET mutants. **(A)** Schematic of designed mutants. **(B)** FPLC chromatograms of SBD1^A^ and SBD2^C^ as isolated protein and tandem. Unlabelled, Cys-containing mutants were loaded on a Superdex 200 10/300 GL column. Tandem SBDs elute around 15.5 ml and are discernible from isolated SBDS, that elute around 17.5 ml. **(C)** SDS Page of fraction A10-B9.

### Section S2. Supplemental Tables

**Supplemental Table S1.**
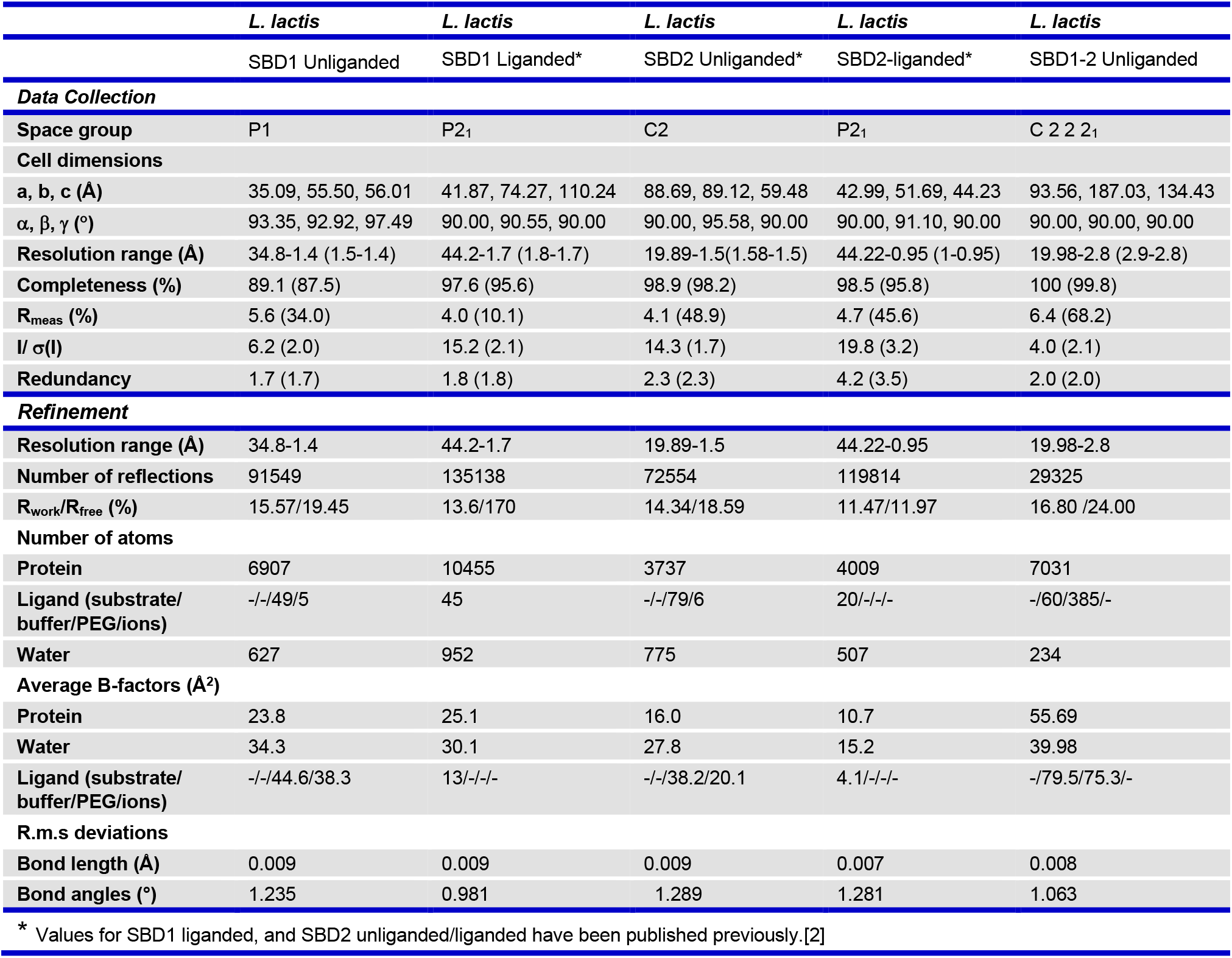
Crystallographic data and refinement statistics. Values in parentheses are for the highest resolution shell.

**Supplemental Table S2.**
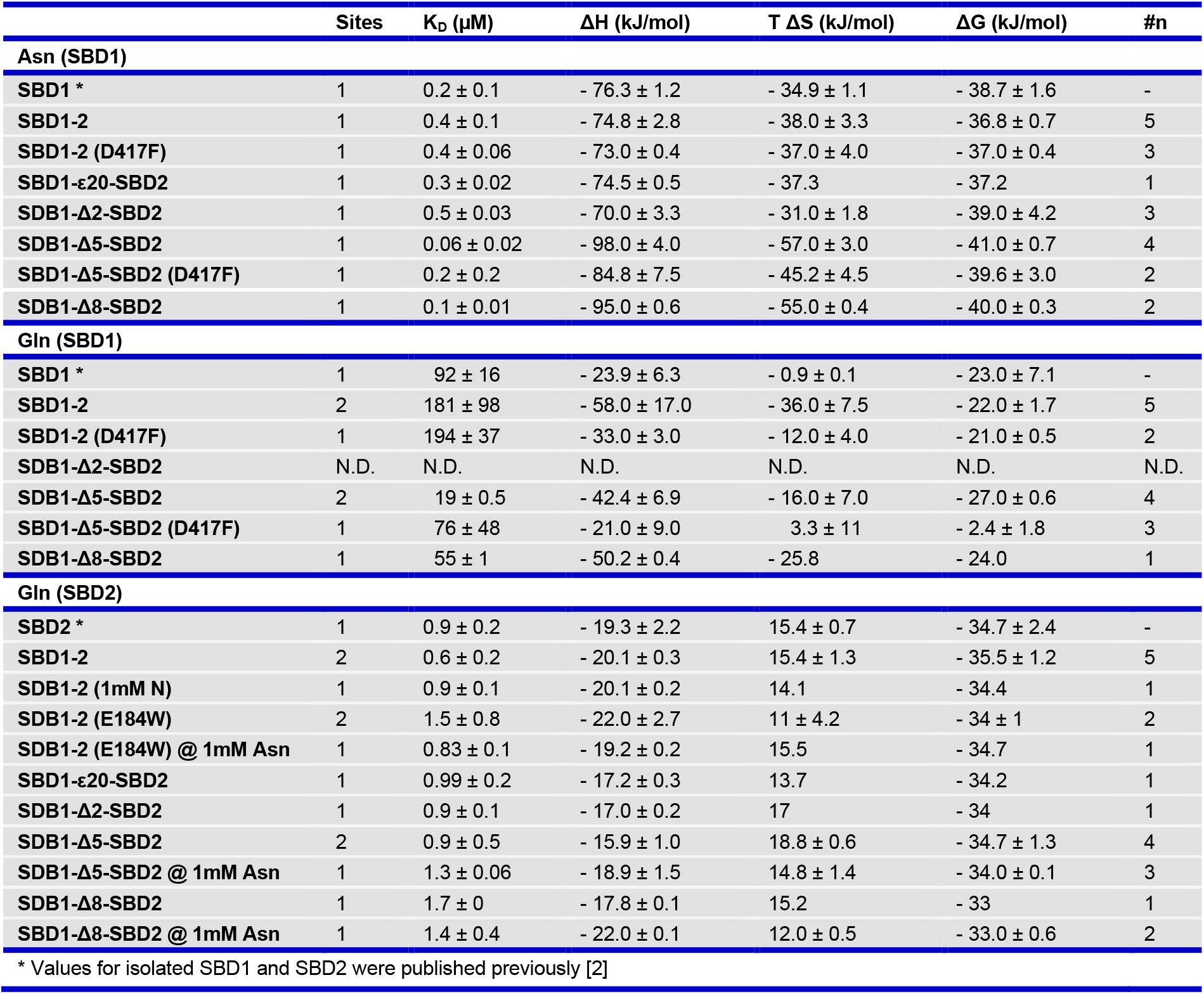
ITC measurements of SBD tandem mutants. Binding affinities of GlnPQ SBD1-2 wild-type and mutants determined by ITC. Error bars represent standard deviations of *n* replicas. N.D. = Not determined. In case of single experiments, *i.e. n* = 1, the provided error margins result from the evaluating fit to the data.

**Supplemental Table S3.** Summary of short notations of proteins derived from SBD1-2. List of all shortened names for all isolated SBDs and tandem mutants. Superscripts refer to the introduced double-cysteine mutants for FRET and PIFE-FRET studies, whereas point mutations are given in brackets. Deletions and extensions of amino acids in the linker sequence are denoted via “Δ” and “ε”, between both SBDs including the number of altered amino acids.

| Unlabelled Proteins |  |  |
| --- | --- | --- |
| Short | Long |  |
| SBDs and Tandem |  |  |
| SBD1 | SBD1 |  |
| SBD2 | SBD2 |  |
| SBD1-2 | SBD1 - SBD2 |  |
| Point Mutations |  |  |
| SBD1 (E184W) | SBD1 (E184W) |  |
| SBD2 (D417F) | SBD2 (D417F) |  |
| SBD1 (E184W)-2 | SBD1 (E184W)-SBD2 |  |
| SBD1-2 (D417F) | SBD1 - SBD2 (D417F) |  |
| Linker |  |  |
| SBD1-Δ2-SBD2 | SBD1 - (ΔT252-K253) - SBD2 |  |
| SBD1-Δ5-SBD2 | SBD1 - (ΔT252-T256) - SBD2 |  |
| SBD1-Δ8-SBD2 | SBD1 - (ΔT252-K259) - SBD2 |  |
| SBD1-ε20-SBD2 | SBD1 - (A251-ggsgggsgggsgggsaaq -T252) - SBD2 |  |
| Mutations |  |  |
| SBD1-Δ5-SBD2 (D417F) | SBD1 - (ΔT252-T256) - SBD2 (D417F) |  |
| Cysteine mutants |  |  |
| Short | Long |  |
| FRET: SBDs and Tandem |  |  |
| SBD1<sup>A</sup> | SBD1 (G87C/T159C) |  |
| SBD1<sup>A</sup>-2 | SBD1 (G87C/T159C) - SBD2 |  |
| SBD2<sup>C</sup> | SBD2 (T369C/S451C) |  |
| SBD1-2<sup>C</sup> | SBD1 - SBD2 (T369C/S451C) |  |
| FRET: Inter-domain distance |  |  |
| SBD1<sup>T</sup>-2<sup>T</sup> | SBD1 (A136C) - SBD2 (T369C) |  |
| PIFE-FRET mutants: SBD1 |  |  |
| SBD1<sup>B</sup> | SBD1 (A136C/S221C) |  |
| SBD1<sup>B</sup>-2 | SBD1 (A136C/S221C) - SBD2 |  |
| SBD1<sup>B</sup>-Δ2-SBD2 | SBD1 (A136C/S221C) - (ΔT252-K253) - SBD2 |  |
| SBD1<sup>B</sup>-Δ5-SBD2 | SBD1 (A136C/S221C) - (ΔT252-T256) - SBD2 |  |
| SBD1<sup>B</sup>-Δ8-SBD2 | SBD1 (A136C/S221C) - (ΔT252-K259) - SBD2 |  |
| SBD1<sup>B</sup>-ε20-SBD2 | SBD1 (A136C/S221C) - (A251-ggsgggsgggsgggsaaq -T252) - SBD2 |  |
| PIFE-FRET mutants: SBD2 |  |  |
| SBD1-2<sup>C</sup> | SBD1 - SBD2 (T369C/S451C) |  |
| SBD1-Δ2-SBD2<sup>C</sup> | SBD1 - (ΔT252-K253) - SBD2 (T369C/S451C) |  |
| SBD1-Δ5-SBD2<sup>C</sup> | SBD1 - (ΔT252-T256) - SBD2 (T369C/S451C) |  |
| SBD1-Δ8-SBD2<sup>C</sup> | SBD1 - (ΔT252-K259) - SBD2 (T369C/S451C) |  |
| SBD1-ε20-SBD2<sup>C</sup> | SBD1 - (A251-ggsgggsgggsgggsaaq -T252) - SBD2 (T369C/S451C) |  |
| Symbol | Domain | Cysteine mutation |
| A | SBD1 | G87C/T159C |
| B | SBD1 | A136C/S221C |
| C | SBD2 | T369C/S451C |
| T | SBD1-2 | A136C/S451C |

**Supplemental Table S4.** Fluorescence anisotropies. Summary of anisotropy values derived from ensemble experiments (Methods section). The anisotropy values for the SBDs are comparable to those of a DNA reference, which confirms that FRET efficiency E reports on inter-probe distances. N.D. = Not determined.

| Data Collection | Anisotropy R |  |  |
| --- | --- | --- | --- |
|  | Cy3 | Cy3B | ATTO647N |
| **Free Dye** | 0.15 ± 0.01 | 0.06 ± 0.01 | 0.03 ± 0.01 |
| **45mer dsDNA / 18 bp** | 0.21 ± 0.01 | 0.13 ± 0.02 | 0.14 ± 0.02 |
| **SBD2<sup>C</sup>** | N.D. | 0.20 ± 0.01 | 0.14 ± 0.03 |
| **SBD1-2<sup>C</sup>** | 0.25 ± 0.01 | 0.16 ± 0.02 | 0.13 ± 0.03 |
| **SBD1-Δ2-SBD2<sup>C</sup>** | N.D. | N.D. | N.D. |
| **SBD1-Δ5-SBD2<sup>C</sup>** | 0.25 ± 0.01 | 0.17 ± 0.01 | 0.19 ± 0.02 |
| **SBD1-ε20-SBD2<sup>C</sup>** | N.D. | 0.15 ± 0.01 | 0.13 ± 0.02 |
| **SBD1<sup>B</sup>** | 0.22 ± 0.03 | 0.19 ± 0.05 | 0.13 ± 0.03 |
| **SBD1<sup>B</sup>-2** | 0.24 ± 0.02 | 0.20 ± 0.03 | 0.14 ± 0.02 |
| **SBD1<sup>B</sup>-Δ5-SBD2** | N.D. | N.D. | N.D. |

**Supplemental Table S4. Fluorescence anisotropies.**
| Data Collection | Anisotropy R |  |
| --- | --- | --- |
|  | Alexa Fluor 555 | Alexa Fluor 647 |
| **Free Dye** | 0.25 ± 0.01 | 0.20 ± 0.01 |
| **SBD2<sup>C</sup>** | 0.26 ± 0.01 | 0.20 ± 0.01 |
| **SBD1-2<sup>C</sup>** | N.D. | N.D. |
| **SBD1<sup>A</sup>** | 0.27 ± 0.01 | 0.19 ± 0.01 |
| **SBD1<sup>A</sup>-2** | N.D. | N.D. |

**Supplemental Table S5.**
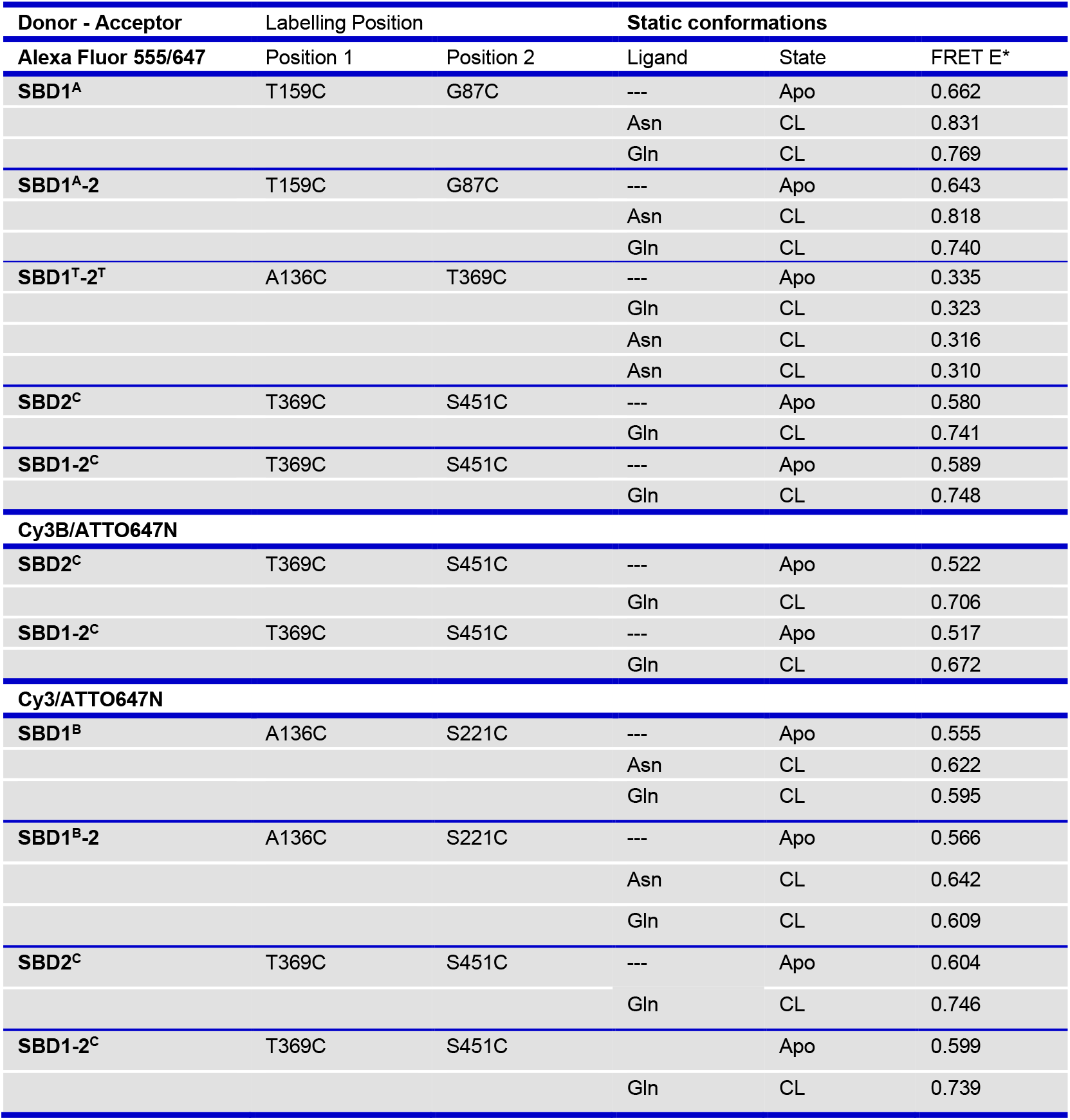
FRET values of isolated SBDs and tandem mutants. Apparent FRET values of isolated SBDs and tandem mutants. FRET values determined via ALEX-spectroscopy of isolated SBD1-(blue) and SBD2-associated (orange) tandem mutants for three fluorophore pairs (Alexa Fluor 555/647, Cy3/ATTO647N and Cy3B/ATTO647N) and various substrates.

| Donor - Acceptor | Labelling Position |  | Static conformations |  |  |
| --- | --- | --- | --- | --- | --- |
| Alexa Fluor 555/647 | Position 1 | Position 2 | Ligand | State | FRET E* |
| SBD1<sup>A</sup> | T159C | G87C | --- | Apo | 0.662 |
|  |  |  | Asn | CL | 0.831 |
|  |  |  | Gln | CL | 0.769 |
| SBD1<sup>A-2</sup> | T159C | G87C | --- | Apo | 0.643 |
|  |  |  | Asn | CL | 0.818 |
|  |  |  | Gln | CL | 0.740 |
| SBD1<sup>T-2<sup>T</sup></sup> | A136C | T369C | --- | Apo | 0.335 |
|  |  |  | Gln | CL | 0.323 |
|  |  |  | Asn | CL | 0.316 |
|  |  |  | Asn | CL | 0.310 |
| SBD2<sup>C</sup> | T369C | S451C | --- | Apo | 0.580 |
|  |  |  | Gln | CL | 0.741 |
| SBD1-2<sup>C</sup> | T369C | S451C | --- | Apo | 0.589 |
|  |  |  | Gln | CL | 0.748 |
| Cy3B/ATTO647N |  |  |  |  |  |
| SBD2<sup>C</sup> | T369C | S451C | --- | Apo | 0.522 |
|  |  |  | Gln | CL | 0.706 |
| SBD1-2<sup>C</sup> | T369C | S451C | --- | Apo | 0.517 |
|  |  |  | Gln | CL | 0.672 |
| Cy3/ATTO647N |  |  |  |  |  |
| SBD1<sup>B</sup> | A136C | S221C | --- | Apo | 0.555 |
|  |  |  | Asn | CL | 0.622 |
|  |  |  | Gln | CL | 0.595 |
| SBD1<sup>B-2</sup> | A136C | S221C | --- | Apo | 0.566 |
|  |  |  | Asn | CL | 0.642 |
|  |  |  | Gln | CL | 0.609 |
| SBD2<sup>C</sup> | T369C | S451C | --- | Apo | 0.604 |
|  |  |  | Gln | CL | 0.746 |
| SBD1-2<sup>C</sup> | T369C | S451C |  | Apo | 0.599 |
|  |  |  | Gln | CL | 0.739 |

**Supplemental Table S6.** Dissociations constants K_D_ of SBDs determined by μs-ALEX. Ligand dissociation constants determined by μs-ALEX. Apparent binding affinities determined via ALEX on double-labeled, tandem mutants. N.D.: not determined.

| *Data Collection* | **Ligand** | **Apparent $K_D$ (μM)** | **Donor** | **Acceptor** | **WT (μM) <sup>3</sup>** |
| --- | --- | --- | --- | --- | --- |
| **SBD1** |  |  |  |  |  |
| **SBD1<sup>B</sup>** | Asn <sup>1</sup> | 0.45 ± 0.04 | Cy3B | ATTO647N | 0.2 ± 0.0 |
| **SBD1<sup>A</sup>** | Asn <sup>1</sup> | 0.35 ± 0.02 | Cy3B | ATTO647N | 0.2 ± 0.0 |
| | Asn <sup>2</sup> | 0.34 ± 0.03 | Alexa Fluor 647 | Alexa Fluor 647 | 0.2 ± 0.0 |
| | Asn | 0.14 ± 0.01 | Alexa Fluor 555 | Alexa Fluor 647 | 0.2 ± 0.0 |
| | Gln <sup>1</sup> | 160 ± 40 | Cy3B | ATTO647N | 92 ± 16 |
| | Gln | 130 ± 10 | Alexa Fluor 555 | Alexa Fluor 647 | 92 ± 16 |
| | His | 120 ± 10 | Alexa Fluor 555 | Alexa Fluor 647 | N.D. |
| **SBD1<sup>A-2</sup>** | Asn | 0.075 ± 0.010 | Alexa Fluor 555 | Alexa Fluor 647 | 0.4 ± 0.0 |
| | Gln | 300 ± 50 | Alexa Fluor 555 | Alexa Fluor 647 | 181 ± 98 |
| | His | 290 ± 70 | Alexa Fluor 555 | Alexa Fluor 647 | N.D. |
| **SBD2** |  |  |  |  |  |
| **SBD2<sup>C</sup>** | Gln <sup>1</sup> | 1.1 ± 0.1 | Cy3B | ATTO647N | 0.9 ± 0.1 |
| **SBD2<sup>C</sup>** | Gln | 1.5 ± 0.3 | Alexa Fluor 555 | Alexa Fluor 647 | 0.9 ± 0.1 |
| **SBD1-2<sup>C</sup>** | Gln | 0.9 ± 0.1 | Alexa Fluor 555 | Alexa Fluor 647 | 0.6 ± 0.2 |
| **SBD1-Δ5-SBD2<sup>C</sup>** | Gln | 1.6 ± 0.1 | Cy3 | ATTO647N | 0.9 ± 0.5 |

<sup>1</sup> Apparent $K_D$ values for Cy3B/ATTO647N have been published previously [3].
<sup>2</sup> Apparent $K_D$ values for Alexa Fluor 555/647 have been published previously [4].
<sup>3</sup> $K_D$ values are determined for unlabeled protein via ITC (see Table S2).

**Supplemental Table S7.** DSC measurements of SBD tandem mutants Thermostability of GlnPQ SBD1-2 wild-type data and mutants determined with DSC. Proteins were added ad 4 μM and 5 mM glutamine and/or asparagine. N.D. = Not determined.

|  | Apo (°C) | + Asn (°C) | + Gln (°C) | + N + Q (°C) |
| --- | --- | --- | --- | --- |
| *Isolated SBD* |  |  |  |  |
| **SBD1 \*** | 60.8 | 72.8 | 64.2 | N.D. |
| **SBD2 \*** | 61.9 | N.D. | 76.2 | N.D. |
| *Tandem-SBD* |  |  |  |  |
| **SBD1-2** | 59.4 | 62.4 / 69.4 | 62.7 / 75.4 | 70.8 / 76.3 |
| **SDB1-Δ2-SBD2** | 59.4 | 61.9 / 69.7 | 62.6 / 74.8 | N.D. |
| **SDB1-Δ5-SBD2** | 58.8 | 61.2 / 70.5 | 65.6 / 73.3 | 73.8 |
| **SBD1-Δ5-SBD2 (D417F)** | 59.3 | 60.7 / 69.3 | 60.7 | 60.7 / 69.3 |
| **SDB1-Δ8-SBD2** | 57.3 | 56.6 / 70.4 | 64.2 | 72.0 |
| * Values involving glutamine been published previously [5] |  |  |  |  |

**Supplemental Table S8.** Fit values of PIFE-FRET mutants. Apparent FRET and Stoichiometry values of isolated SBDs and tandem mutants. Summary of apparent FRET and Stoichiometry values for all mutants in the *apo-* and closed-liganded state that are employed within the PIFE-FRET assay. For mutants with respect to SBD1^A^, i.e., SBD1(A136/S221), that show no PIFE, only Cy3/ATTO647N labelling has been carried out. For mutants with respect to SBD2^C^, i.e., SBD2(T369/S451), labelling with Cy3 and Cy3B/ATTO647N has been carried out. For saturating concentrations of the ligand, either 200 μM asparagine or 1 mM glutamine have been employed. Values represent the mean and standard deviation of n > 2 replica in the *apo-* (A) and closed-liganded (CL) state.

| PIFE-FRET assay | FRET E* |  | Stoichiometry S* |  | #n |
| --- | --- | --- | --- | --- | --- |
| Mutants | Apo | CL | Apo | CL | A/CL |
| **Cy3** |  |  |  |  |  |
| SBD1<sup>B</sup> | 0.566 ± 0.017 | 0.622 ± 0.039 | 0.284 ± 0.012 | 0.320 ± 0.026 | 11/4 |
| SBD1<sup>B</sup>-2 | 0.555 ± 0.015 | 0.641 ± 0.031 | 0.289 ± 0.017 | 0.324 ± 0.024 | 12/3 |
| SBD1<sup>B</sup>-Δ2-SBD2 | 0.592 ± 0.007 | 0.617 ± 0.009 | 0.287 ± 0.005 | 0.295 ± 0.009 | 6/2 |
| SBD1<sup>B</sup>-Δ5-SBD2 | 0.566 ± 0.005 | 0.625 ± 0.006 | 0.290 ± 0.009 | 0.319 ± 0.006 | 4/2 |
| sSBD1<sup>B</sup>-Δ8-SBD2 | 0.558 ± 0.003 | 0.628 ± 0.005 | 0.286 ± 0.007 | 0.318 ± 0.006 | 5/2 |
| SBD1<sup>B</sup>-ε20-SBD2 | 0.566 ± 0.012 | 0.621 ± 0.013 | 0.286 ± 0.016 | 0.323 ± 0.013 | 8/2 |
| SBD2<sup>C</sup> | 0.613 ± 0.018 | 0.730 ± 0.016 | 0.291 ± 0.007 | 0.369 ± 0.007 | 13/3 |
| SBD1-2<sup>C</sup> | 0.616 ± 0.021 | 0.736 ± 0.010 | 0.306 ± 0.016 | 0.371 ± 0.018 | 14/3 |
| SBD1-Δ2-SBD2<sup>C</sup> | 0.575 ± 0.014 | 0.741 ± 0.013 | 0.312 ± 0.006 | 0.382 ± 0.012 | 5/2 |
| SBD1-Δ5-SBD2<sup>C</sup> | 0.569 ± 0.014 | 0.722 ± 0.009 | 0.339 ± 0.007 | 0.400 ± 0.002 | 6/2 |
| SBD1-Δ8-SBD2<sup>C</sup> | 0.578 ± 0.005 | 0.689 ± 0.007 | 0.268 ± 0.012 | 0.333 ± 0.008 | 7/2 |
| SBD1-ε20-SBD2<sup>C</sup> | 0.634 ± 0.028 | 0.779 ± 0.025 | 0.300 ± 0.107 | 0.370 ± 0.018 | 5/2 |
| **Cy3B** |  |  |  |  |  |
| SBD2<sup>C</sup> | 0.517 ± 0.016 | 0.688 ± 0.025 | 0.590 ± 0.012 | 0.566 ± 0.013 | 4/2 |
| SBD1-2<sup>C</sup> | 0.517 ± 0.012 | 0.679 ± 0.014 | 0.594 ± 0.007 | 0.569 ± 0.016 | 14/3 |
| SBD1-Δ5-SBD2<sup>C</sup> | 0.447 ± 0.011 | 0.619 ± 0.019 | 0.609 ± 0.012 | 0.579 ± 0.011 | 3/2 |
| SBD1-ε20-SBD2<sup>C</sup> | 0.514 ± 0.031 | 0.688 ± 0.039 | 0.595 ± 0.008 | 0.568 ± 0.011 | 3/2 |

